# Microbial communities do not drive chronic *Quercus* spp. decline but exacerbate tree weakening: a multi-kingdom soil–root continuum study in the Chantilly forest

**DOI:** 10.64898/2026.11.17.766935

**Authors:** A. Deveau, L. Eichenlaub, L. Gigleux, F. Guinet, L. Saint-André, M. Gómez Gallego, C. Robin

## Abstract

In European forests, several species of *Quercus* spp. are currently subject to major declines, defined as a slow deterioration of the tree caused by the interaction of abiotic and biotic factors, in which drought often plays an important role. The decline of holobionts, like oak trees, may have consequences that extend beyond the tree physiology. To better understand the interactions between chronic oak decline and oak associated microbial communities and their functional consequences, we analysed the bacterial, fungal and oomycetes communities in the bulk soil, the rhizosphere and the roots of pairs of declining and non-declining oaks in the Chantilly forest (France) by a metabarcoding approach. Chosen pairs of oaks were distributed on contrasted soil types to determine the influence of soil properties on the tree-microbe interactions during decline. Rhizospheric soil microbial enzymatic activities were also measured to assess the effect of oak decline on microbial-driven nutrient cycling activities. We observed a differential response of microbial communities to decline as a function of the kingdom. Bacteria were more sensitive to oak decline, particularly in the bulk soil and on podzol soil. By contrast, the composition of fungal communities was overall little impacted by oak decline, no matter the soil type, but their functional abilities may be partly impaired. Laccase and chitinase activities were reduced in the rhizospheric soil of declining oaks growing on brunisol and luvisol. Furthermore, the proportion of ectomycorrhizal fungi with high biomass exploration types were reduced by half in the roots of declining trees while the pathogenic *Phytophthora cactorum* and *P. plurivora* were more frequent. Altogether, our data suggest that chronic oak decline affects directly and indirectly the composition and functional abilities of oak associated microbial communities in a soil dependent manner.

## INTRODUCTION

Forest ecosystems face both abiotic and biotic disturbances, which affect trees based on their vulnerability and exposure levels and whose frequency and intensity are rising with the acceleration of climate change (Trumbore et al., 2015; Seidl et al., 2017). Their co-occurrence or succession over time often triggers a process of decline in trees, defined as a slow and progressive deterioration in metabolism and functioning, resulting in crown decline, and which can lead to death. Characteristic symptoms of decline (mainly yellowing and wilting of leaves, defoliation and branch mortality) are induced by predisposing, inciting or aggravating factors, all depending on the local or regional context (Delatour, 1983, Manion, 1981). Predisposing factors act over the long term to weaken the trees, while inciting factors are short term stresses that trigger the decline, and contributing factors, mostly opportunistic organisms, act on the weakened trees to increase or to speed up the level of decline and mortality.

Numerous cases of oak decline were reported as early as the 18th century in Europe and then throughout the world (Gossling et al. 2024). The decline of *Quercus petraea* (Mattuscka) Liebl. and *Q. robur* L. has been a chronic problem in Europe for the past 40 years. Moreover, a complex disease, called Acute Oak Decline (AOD), is currently affecting oak forests in several European countries. Mediated by abiotic factors (temperature and precipitation), AOD would be triggered by an insect, *Agrilus biguttatus*, and a complex of bacterial species (Denman et al. 2014).

The most commonly cited causal factors of chronic (as opposed to acute) oak decline are drought, frost, silvicultural practices, insects or fungal pathogens, although other hazards may have a strong effect (Thomas et al., 2003). Different fungal pathogens may act as predisposing, inciting or aggravating factors of oak decline. Among them, root pathogens worsen the host’s health status, by degrading fine roots or main roots and collar. For instance, butt and root rot pathogens may be either predisposing (for example *Collybia fusipes*, Marçais & Caël, 2000) or aggravating (for example *Armillaria* species, Marçais & Bréda 2006). Several species of *Phytophthora* have also been suggested as key factors in oak decline. Rising temperatures and extreme rainfall events leading to waterlogging favour many of these oomycetes, because active root infection from soil inoculum depends on free soil water that enables zoospore movement toward host roots (Judelson and Blanco, 2005). Moreover, trees whose root system has been partially destroyed by *Phytophthora* infection, may be more vulnerable to drought (e.g. Jönsson et al. 2003b). However, while numerous studies have corroborated the involvement of *P. cinnamomi* in the decline of Mediterranean oaks, it is more difficult to generalise the role of *Phytophthora* spp. in the decline of deciduous oaks. While *P. quercina* as been frequently associated with declining oaks (*Q. robur*, *Q. petraea*, *Q. cerris* L., Q. *pubescens* Willd. and *Q. ilex*) around Europe (Jung et al., 1999; Jönsson et al. 2003a, Cooke et al., 2005), its pathogenicity on oak seedlings appeared dependent on climate and soil type (Jönsson et al. 2003b, Jönsson, 2004, Jönsson-Belyazio & Rosengren 2006). Other *Phytophthora* species have been reported as pathogenic to oak, causing significant root death, and associated with oak decline, including *P. plurivora*, *P. cambivora* and *P. cactorum* (Tkaczyk et al. 2020, Jankowiak et al. 2014).

Independently of pathogens, roots are naturally massively colonised by complex microbial communities, dominated by bacteria and fungi, which play a key role in the nutrition and health of trees (Uroz et al. 2016). Among the fungi, ectomycorrhizal (ECM) fungi are key players by providing protection to the roots against pathogens and drought while gathering soil nutrients and water for their tree hosts in exchange of photosynthetic carbon (Smith & Read 2007). Each tree root system is generally colonized by a few dozen ECM species which may differ in their functional abilities in terms of soil exploration and resource acquisition capabilities (Courty et al. 2007), and in terms of carbon cost for the host tree (i.e. high vs low biomass ECM fungi, Hobby & Agerer 2010). In addition to symbiotic ECM fungi, saprophytic fungi that colonise dead organic matter and root rhizospheres play a crucial role in releasing nutrients from organic matter. As such, they regulate nutrient cycling and carbon sequestration in soils (Baldrian 2017). Bacteria are also central to tree nutrition and health by contributing to the mobilisation of nutrients, promoting growth and interfering with pathogen development (Lladó et al. 2017). They also play a key role in the nitrogen cycle in soil through nitrogen fixation, nitrification and denitrification. Lastly, on top of their functional roles, soil communities also serve as a reservoir of microorganisms for the colonisation of root systems (Fracchia et al. 2021, 2024). The position of fungi and bacteria at the interface between tree roots and soil results in a sensitivity not only to changes in soil chemistry and edaphic conditions but also to changes in tree carbon allocation through both root exudation and nutrient trade with ECM (Baldrian et al. 2023). Thus, impacts on soil microbial communities are likely to have a detrimental effect on tree performance. Conversely, a poor tree physiological condition is likely to have an impact on the microbial communities associated with its roots. The intensity of this effect is expected to be stronger in the roots (endosphere) and its vicinity (rhizosphere) than in the bulk soil which is not under the direct influence of root exudates (Arnault et al. 2023). Conversely, bulk soil is expected to be more susceptible to the direct effects of abiotic stresses such as drought or heat (Arnault et al. 2023).

With regard to oak decline, which occurs in a context of climate change characterised by increased frequency and intensity of droughts and heat waves, two questions arise: to what extent is the soil microbiota involved in the decline process and, conversely, does oak decline affect the composition and functional activities of the forest soil microbiota? Such questions have been addressed in Mediterranean forests of *Q. ilex* and *Q. suber* experiencing decline (Montecchio et al. 2004, Corcobado et al. 2014, García-Angulo et al., 2020, Gómez-Aparicio et al., 2022), or for AOD (Corcobado et al. 2014, Barsoum et al. 2021, Pinho et al. 2020, Gómez-Aparicio et al., 2022). Surprisingly, only four studies on *Q. robur* and *Q. petraea* exist, despite both species being prevalent in European forests (Causin et al. 1996, Kovacs et al. 2000, Scarlett et al. 2021, Lopez-Garcia et al. 2025). While Causin and colleagues and Kovacs and colleagues concentrated solely on ECM species and Scarlett and colleagues on nitrogen-cycle-related bacteria, Lopez-Garcia and colleagues adopted a broader approach combining metabarcoding and baiting techniques to compare fungal, bacterial, and *Phytophthora* communities in the rhizosphere of ten healthy versus ten declining oaks in a southern Sweden forest. Interestingly, their results suggest that microbial communities vary with tree health and that fungal and bacterial communities react differently. In addition, they detected *Phytophthora* pathogens in the rhizosphere of numerous seemingly healthy trees, suggesting that these trees may be asymptomatic carriers (Lopez-Garcia et al. 2025). These results raise the question whether a dysbiosis of forest soil microbiota could be a predisposing factor of decline, following the conceptual model of Manion (1981). The second key question arising from the work of Lopez-Garcia concerns the extent to which their results can be extrapolated to other European oak forests. Indeed, oak stands occur across a wide range of climates—from oceanic to continental—as well as on diverse soil types (Eaton et al. 2016), both recognized as primary drivers of microbial diversity and community composition (Baldrian 2023).

Chantilly forest provides an interesting field for the study of the link between soil microbiota and chronic oak decline. This forest is dominated by *Q. robur*, *Q. petraea* and a few hybrids between these two species (B. Bracchi, personal communication), growing on a mosaic of soil types, which is well representative of dominant soils found in France. Since 2010, recurrent droughts have been associated with an exacerbated decline of oaks which were showing a loss of vitality, without any indication of AOD. To clarify the relationships between microbiota from different compartments (soil, rhizosphere and roots) and chronic oak decline, this study was set out to test the following hypotheses: (i) microbiota of healthy oaks in a forest under chronic decline show signs of dysbiosis, (ii) oak decline effects on microbiota composition and diversity depend on taxon or kingdom, soil type, soil-root continuum and functional guild, (iii) the alteration of root and rhizospheric microbial communities in declining oak reduces microbial functional abilities in terms of nutrient cycling. To test these hypotheses, we used pairs of oaks of contrasting health status sampled in the Chantilly forest. A culture-independent approach relying on ITS and 16S rRNA metabarcoding was performed to study fungal, bacterial and oomycete communities in three compartments (soil, rhizosphere and roots) and enzymatic activity were measured on rhizospheric soil samples to assess functional abilities in terms of nutrient cycling as a function of soil type and health status.

## MATERIALS AND METHODS

### Study site and sampling

The study was conducted in the Chantilly forest, 40 km North East of Paris (France, **Figure 1**). and dominated by *Q. robur*, *Q. petraea* (and a few hybrids between these two species, B. Brachi, personal communication). The Chantilly Forest is an ageing forest growing on soil with reduced water capacity well representative of dominant soils found in France. Since 2010, this forest has suffered recurrent droughts and intense summer heat waves, compounded by attacks from cockchafers. This combination of factors can weaken trees and even cause them to die. In 2020, 23% of oaks of Chantilly forest were classified as declining and 16% as being in an advanced state of decline.

**Figure 1.**
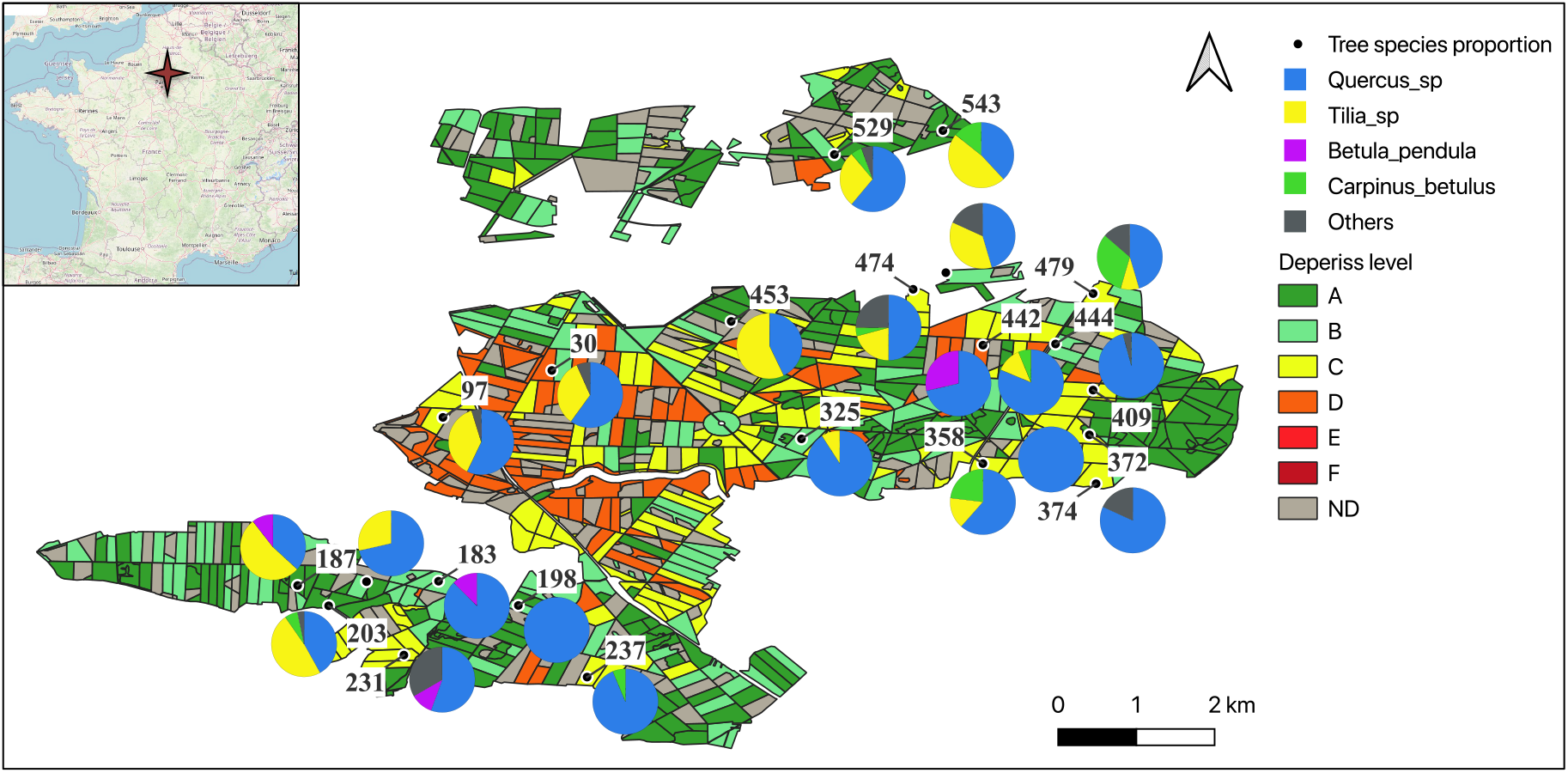
Localisation and properties of study plots in the Chantilly forest. Decline level, as determined through DEPERIS method, is depicted for the entire Chantilly forest from no mortality (green) to high mortality (red). Piecharts indicate the proportion of tree indivudal per species in each sampled plot.

The study site is made of 22 permanent monitoring plots set up in 2021 (**Figure 1, Suppl. Table 1)**. Each plot was characterized in depth in terms of soil physico-chemical properties, vegetation, tree genetics and tree health (see complementary site description in **supplementary information**). All data regarding plot characteristics are available on the French dataverse (Saint-André et al. 2024, b, Santenoise et al. 2024, Sénéchal-Chevalier et al. 2024). Crown deterioration measurements were made on all oaks of the 22 plots in 2021 using the DEPERIS scoring method which comprises six rating classes (https://agriculture.gouv.fr/la-methode-deperis-comment-quantifier-et-mesurer-letat-de-sante-dune-foret-et-son-evolution, cf. **supplementary information**). For our study, we defined two tree categories: trees with more than 50% of branch mortality and more than 50% of branching loss in the crown part not affected by mortality were referred to as declining (D) trees whereas other trees were considered as non-declining trees (hereafter referred to as ND). No characteristic symptoms of AOD were detected on tree trunks during this assessment. We selected 29 pairs of oaks (one D and one ND tree) located on the same plot with a similar circumference at 1.30 m height (+/-9 cm). The distance between the trees of each pair ranged from 10 to 45 m. The 22 plots were dominated by *Q. robur* and *Q. petraea:* 14 plots were composed of more than 50% of oak trees and two were monospecific. Five plots were composed of more than 33% of *Tilia* sp. Other tree species were *Betula pendula* and *Carpinus betulus* (**Figure 1**). The 29 pairs of trees were spread across four soil types (**Suppl. Table 1**). Soil types were grouped together based on their physico-chemical characteristics as follows: calcisol and rendzina formed one group (CA-REN, 7 pairs), as did luvisol and luvisol-redoxisol (LUV, 6 pairs). Dystric brunisol (DYB, 8 pairs) and podzol (POD, 8 pairs) were considered separately. The plots were thus evenly distributed among four groups of soil types with contrasting water pH intervals (27% of the plots were on DYB [pH: 3.9-5.4]; 23% on CA-REN [pH: 5.2-7.6]; 23% on LUV [pH: 3.9-5.1]; 27% on POD [pH: 4.0-4.7]). Three humus categories were identified (dismull, mesomull, oligomull in 10%, 38 and 52% of the plot, respectively **Suppl. Table 1**).

For each tree of each pair, roots, rhizospheric soil and bulk soil were sampled at one meter of the trunk to a depth of ca. 15 cm. After litter removal with a rack, three cubes of soil (15 x 15 × 10 cm) were sampled with a shovel at three cardinal points (N, S, E, W depending on field configuration). Cubes were gently broken to extract root systems without removing the rhizospheric adherent soil. Collected materials from each compartment taken at the three cardinal points were pooled together to form one sample type per compartment and per tree. The roots identified as oak roots based on their appearance (colour, size, shape) were sorted and collected to form the root sample. Rhizospheric soil (Rh) was collected by shaking and rubbing with hands the roots in plastic bags. A handful of roots devoid of adherent soil particles were further collected to form the “Root” (R) sample. Bulk soil (BS) samples, passed through a 2-mm sieve, were made up of soil free from any root system and not under the influence of root activity. All materials used to collect and process samples (shovels, sieves) were cleaned with water and with 70% ethanol between each tree to avoid DNA cross contaminations. All samples were frozen at -20 °C after reaching the laboratory before further use, except a subset of rhizospheric soil which was kept at 4 °C for enzymatic activity measurements.

### DNA extraction and sequencing library constructions

DNA was extracted with the Dneasy Powersoil pro kit (Qiagen, Hilden, Germany) from 250 mg of soil and rhizosphere samples and from 50 mg with DNeasy Plant mini kit (Qiagen, Hilden, Germany) for root samples. All DNA extractions were performed in three replicates. DNA concentrations and quality were measured with NanoDrop spectrophotometer (Thermo Fisher Scientific, USA). The three sample repetitions were pooled with equimolarity and the final concentration diluted at 10 ng/µl.

A two-step PCR approach was chosen to barcode tag templates. Three libraries were made to specifically target three taxa: oomycetes, bacteria and fungal taxa. Negative controls during the PCR process have been made replacing the DNA by sterile Milli-Q® water and mock communities of known fungal, oomycetes and bacterial compositions were added as quality controls. Fungal amplicons were generated at ∼250 bp sequences by amplifying the ITS1 region. Primers ITS1F [2 µM] (5’ CTTGGTCATTTAGAGGAAGTAA 3’) and the ITS2 [2 µM] (5’ GCTGCGTTCTTCATCGATGC 3’) (Pauvert *et al.,* 2019) were used in the 13-µl reaction volume in association with 2.5 µl of extracted DNA, bovine serum albumin (BSA) [10 ng/µl] and Taq 2X KAPA HiFi HotStart Ready Mix enzyme (KAPA Biosystems, Wilmington, MA, USA) completed by sterile Milli-Q® water. PCR conditions were: initial denaturation at 94 °C for 3 min, 35 cycles of denaturation at 94 °C for 45 s, annealing at 53 °C for 30 s, initial extension at 72 °C for 45 s and 1 cycle of final extension at 72 °C for 10 min.

Amplicons of oomycetes in samples, mock and negative controls were generated at ∼250 bp sequence of the internal transcribed spacer (ITS) and 18S region fragment by using a nested PCR (Catala *et al.,* 2015). For the first PCR round, *Phytophthora*-specific primers 18Ph2F [2 µM] (5’ GGATAGACTGTTGCAATTTTCAGT 3’) and 5.8S-1R [2 µM] (5’ GCARRGACTTTCGTCCCYRC 3’) were used, and primers ITS6 [2 µM] (5’ GAAGGTGAAGTCGTAACAAGG3’) and 5.8S-1R [2 µM] (5’ GCARRGACTTTCGTCCCYRC 3’) for the nested PCR. The 12.5 µl of reactional volume were composed of 2.5 µl of extracted DNA (1rst round) or 2.5 µl of PCR1 product (nested PCR), BSA [10 ng/µl] and Taq 2X KAPA HiFi HotStart Ready Mix enzyme (KAPA Biosystems, Wilmington, MA, USA) completed by sterile Milli-Q® water. Amplification conditions were as follows: initial denaturation at 95 °C for 2 min, 30 cycles (1rst round) or 25 cycles (nested round) of denaturation at 95 °C for 20 s, annealing at 60 °C for 30 s, initial extension at 72 °C for 30 s and 1 cycle of final extension at 72 °C for 7 min.

Bacterial amplicon libraries of the 16S rRNA were produced by using 799F (5’ AACMGGATTAGATACCCKG 3’) and 1115R (5’ AGGGTTGCGCTCGTTG 3’) primers to exclude amplification of tree mitochondrial and plastid DNA. PCR were performed for each sample with 2 µL of pooled extracted DNA adjusted at a concentration of 10 ng/µl using 2.5X Phusion flash high fidelity master mix (Thermo Scientific), with 1.5 µl of forward and reverse primer mix (10 µM), BSA (1 mg/ml) 2 µl, DMSO 100% 0.9 µl and completed by sterile Milli-Q® water to a volume of 30 µl. PCR using conditions were the following ones: initial denaturation at 98 °C for 5 s, 30 cycles of denaturation at 98 °C for 5 s, 78 °C for 10 s, 52 °C for 20 , 72 °C 15 s.

For the three libraries, correct amplification of all PCR products was verified by electrophoresis. The clean up of PCR1, PCR2 and the sequencing were performed by the Plateforme Génome Transcriptome de Bordeaux (PGTB - Cestas, France). Each library was sequenced on the MiSeq Instrument (Illumina, San Diego, CA, USA) by making one run with the reagent kit Nano V2 (2 x 250 bp) for the rhizosphere ITS amplicons and two runs with the reagent kit V2 (2 x 250 bp) for the rest. After quality control, FASTQ files of each sample were exported for bioinformatics analysis.

### Bioinformatic treatments

Bioinformatics treatments of the three libraries were performed with the FROGS pipeline (Escudié *et al.,* 2018; Bernard *et al.,* 2021) on the Galaxy interface. For each library independently, sequences were clustered into Amplicon Sequence Variants (ASVs) based on the iterative SWARM algorithm and then chimera were removed. As suggested by Escudié and collaborators (2018), ASVs with a number of reads lower than 5·10^−5^ percent of total abundance were removed. For the fungal library, ASVs sequences matching the ITS region were filtered using the ITSx filter implemented in FROGS. When no assignment to a fungal taxon was possible, they were discarded. Fungal and bacterial sequences were affiliated using the UNITE Fungal Database, v9 (Nilsson *et al.,* 2019), the Silva Database 138.1 and the UNITE Eukaryote Database v9, respectively. Fungal and bacterial multi affiliated ASVs were manually reassigned by using reference sequences in the NCBI Genbank database. Oomycete ASVs were affiliated using the Phytophthora ID (http://phytophthora-id.org/) database.

The average number of fungal and bacterial reads reached 10 583 and 16 327 reads per sample and varied from 243 to 37 797 and from 8 453 and 22 618, respectively. In order to achieve an equal number of reads in all samples, the *rarefy_even_depth* function from the *Phyloseq* package (Mc Murdie & Holmes, 2013) in R (R Core Team, 2020) was used with a depth of 4 500 and 8 453 reads per sample for fungi and bacteria, respectively. A total of 33 samples were then removed, ten from the bulk soil, 19 from the rhizosphere and four from the roots compartment due to low fungal ASV coverage (**Suppl. Table 2**). The rarefied ASV tables were used to assess the relative abundance of fungal and bacterial taxa, as well as α- and β- diversity. FUNGuild (Nguyen *et al.,* 2016) was used to classify each fungal ASV into an ecological trophic guild. A confidence threshold was applied to only keep “highly probable” and “probable” affiliated trophic guilds and the other ASVs were assigned as “unidentified”. Three trophic guild categories were considered: ectomycorrhizal fungi (EMF), plant pathogens, and saprophytes. All other assigned guilds were grouped into “Others”. Mycorrhizal exploration types were assigned to each taxon following Agerer (2001, 2006) and assignments made by Suz *et al*. (2013). They were further classified as “low biomass” (contact, short- and medium-distance smooth exploration types) and “high biomass” (medium-distance fringe, medium-distance mat and long-distance exploration) based on Hobbie & Agerer (2010) and Suz *et al*. (2013).

Four species of *Phytophthora* present in the mock community were not retrieved in this mock sample: no reads were assigned to *P. syringea*, *P. taxon forest soil*, *P. x multiformis* or *Py. sylvaticus.* Only three ASV were detected in the negative controls. Two were assigned to *P. cambivora* and one to *P. alni* subspecies *uniformis*. These two species were included in the mock sample but never detected in the studied samples. The number of oomycete reads per sample varied from 0 to 59 999. A threshold of 300 reads was applied to consider that oomycetes were present in one sample.

### Enzymatic activities

We monitored eight extracellular enzyme activities involved in carbon, nitrogen and phosphorus mobilization involved in the decomposition process of soil organic matter: β-glucosidase, cellobiohydrolase, xylosidase, glucuronidase, N-acetylglucosaminidase, leucine amino-peptidase, acid phosphatase and laccase as described by Pritsch *et al*. (2011). Two grams of sieved rhizospheric soil was added to 20 ml of deionised sterile water to give a final concentration of 1:20 (w/v). The solutions were incubated at room temperature under agitation for two hours. After incubation, 200 μl of the diluted solution was added to 50 μl of substrate solution (methylumbelliferone fluorochrome) in a 96-well plate (Sarstedt, Newton, NC,USA). For laccase activity, 100 µl of the diluted solution was added to 100 µl of 2,2′-Azinobis(3-Ethylbenzthiazoline-6-Sulfonate) (ABTS) substrate solution as described in Maillard *et al*. (2019). After a short period of incubation at 25 °C in the dark, using a horizontal shaker (40 min for β-glucosidase, 60 min for cellobiohydrolase, 40 min for xylosidase, 40 min for glucuronidase, 15 min for acid phosphatase, and 60 min for N-acetylglucosaminidase and leucine aminopeptidase, 120 min for laccase), the plates were centrifuged at 4500 g for 5 min. Then 100 μl of the supernatant was transferred into a black opaque 96-well microplate (Perkin-Elmer, Life Sciences) containing 100 μl of the stopping buffer (Tris 2.5 M pH 10–11), except laccase test for which 200 µl were transferred to a transparent 96-well plate and for which absorbance at 420 nm was immediately measured using Tecan microplate reader (Infinit M200 Pro). For other assays, all fluorometric measurements were carried out using a Tecan microplate reader (Infinit M200 Pro) with an excitation wavelength of 360 nm and an emission wavelength of 450 nm. Gain was adjusted for each substrate to optimize signal/noise ratio as following: 80 for β-glucosidase, 109 for cellobiohydrolase, 86 for xylosidase, 108 for glucuronidase, 65 for acid phosphatase, and 72 for N-acetylglucosaminidase and 59 for leucine amino-peptidase. Standard curves were produced for each value of gain using 5 levels of concentrations of methylumbelliferone to link fluorescence values to pg of substrate produced. Control measurements without soil solution and with soil solution but without substrate were made for each enzymatic assay. Enzymatic activities were expressed in pmol of the substrates released per g of dry soil per min. All chemicals were purchased from Sigma-Aldrich (France).

### Statistical analyses

Statistical analyses and data representation were performed with the R software (R Core Team. 2016) version R-4.2.3 and the RStudio interface (RStudio, Inc). Maps were generated with QGIS v3.40.

#### Analyses of non-declining microbial communities associated with oaks

PERMANOVA (*adonis2* function, *vegan* package, 999 permutations) was used to test the effects of compartment, soil properties, and tree properties on community structures (by = “margin”), using Bray-Curtis dissimilarity measure for fungal and bacterial data and Jaccard dissimilarity measure for oomycetes data. Plot identity was treated as random factors (strata = “argument”).

To test for the effects of compartment, soil properties, and tree characteristics on richness and diversity (Inverse Simpson index) of fungal and bacterial communities of healthy trees, we used a generalized additive mixed model (*gamm* function, *mgvc* package). Soil type, compartment (BS, Rh, R), soil properties (pH, total organic carbon, total organic nitrogen…) and tree characteristics were included as fixed effects. Plot identity was included as a random factor in the model. Poisson and Gamma distributions were used for the richness and the Inverse Simpson models, respectively. Backward selection using AIC was used to obtain the best model. Residual diagnostics were analysed to assess model fit and distribution adequacy. Pairwise comparison between soil types and sample compartments were further performed (*emmeans* function, *emmeans* package) followed by the adjustment of *p* values to account for multiple comparisons (*p.adjust* function, BH method, Benjamini & Hochberg, 1995).

A similar model was used to test the difference in enzymatic activities of rhizospheric soils of non-declining trees depending on the soil type. In this case, each enzymatic activity was used as a variable response in a separate model. We used a Gaussian distribution with identity link, which yielded the best residual diagnostics. Correlations between cellobiohydrolase enzymatic activity and the relative abundances of *Lactarius* the rhizospheric soil samples was calculated using Kendall correlation with the *corr.test* function. To further analyse the relationships between cellobiohydrolase activity, *Lactarius* rarefied abundance and the physico-chemical parameters, tree species, circumference, and stand variables, we performed individual generalised additive mixed models with a *Gaussian* family and identity link and selected best model using AIC as previously described. The approach was further extended to look for correlations between the rarefied abundance of the 12 most abundant ECM taxa and other enzymatic activities, using Kendall correlation. Combinations giving significant correlation (*p*. value < 0.05) were further analysed using generalised additive mixed models as described for cellobiohydrolase as a function of *Lactarius* abundance.

The same approach was used to analyse the abundance of *Phytophthora* species as a function of the rarefied abundance of the 12 most abundant fungal and bacterial taxa, and soil and tree properties.

To analyse the drivers of oomycete presence, we modelled the oomycete detection as a function of compartment and soil type (fixed factors) in non-declining trees (*glmmTM* function, *glmmTMB* package, with tree pair nested within plot as random factors). We then performed pairwise comparison and *p* value correction as described above.

#### Analyses of oak non-declining vs declining trees’ microbial communities

We repeated the PERMANOVA analyses (*adonis2* function, *vegan* package, 999 permutations) for the whole data of each compartment and adding the health status, to test the effects of health status and soil type on community structures (by = “margin”).

To test for the effect of the health status on alpha-diversity, we used a generalized additive mixed model considering health status, soil type and their interaction as fixed effects. The effects of plot and pair, plot alone or pair alone as random factors were tested by comparing AIC values of the three models and tree pair identity was retained as the random-effect component of the model. Poisson and Gamma distributions were used for the richness and the Inverse Simpson models, respectively. Residuals were analysed to validate distribution families. Pairwise comparison between soil types and health status were performed as described above.

As our data were highly dimensional including metabarcoding of three different organisms as well as soil and tree variables, we used the Elastic Net algorithm to identify the parameters that best discriminated between declining and non-declining trees. Elastic Net is a machine learning approach well suited to datasets where the number of explanatory variables greatly exceeds the number of observations. We used 6 365 variables, including: rarefied numbers of reads of each bacterial ASVs present in at least three samples and with a minimal abundance of five reads, bacterial calculated alpha diversity indices (richness, Chao1, Inverse Simpson), relative abundances of fungal trophic guilds (saprotrophs, ECM, pathogens, others, unknown), total oomycete numbers of reads of each oomycete taxon, and enzymatic activities. We first optimized the parameter lambda with the function *cv.glmnet* (R package *glmnet*). As the number of observations was low, the optimisation of the parameter lambda was dependent on the number of folds in which the data base was split in this first step. Therefore, we used a bootstrap approach as follows: we run 50 times the *cv.glmnet* model for each number of folds (*nfolds* argument) between 4 to 15. For each run, we obtained an optimal lambda which we used in the *glmnet* model to obtain the selected variables. We obtained a total of 600 lists of selected variables for each model. We then identified the variables that had been selected in at least 350 out of the 600 total models, as explanatory of the tree health status. Those variables were subsequently validated using generalized additive mixed models. We run individual *gamm* models using each of those variables as a fixed factor, and the health status as a response variable. We then corrected the obtained *p* values by using the *fdr* correction method of the *p.adjust* function.

The effect of the health status on the proportion of different fungal trophic guilds and on the proportion of high vs low biomass ECM fungi was tested using a generalised additive mixed model considering soil type, health status and their interaction as fixed effects with a beta family distribution. Tree pair identity was considered as a random factor in the model. Pairwise comparison between soil types and health status were performed and *p* values adjusted as previously described.

To analyse the relationship between oak decline and the presence of *Phytophthora* species, we performed individual models with the number of reads of a given *Phytophthora* species as a response variable, the tree health status as fixed factor, the logarithm of the total number of reads as an offset, and the sample type and the tree pair nested within plot as random factors. Correlations between the abundance of *Phytophthora* species in declining vs non-declining tree samples and the rarefied abundance of the 12 most abundant fungal and bacterial taxa were first calculated using Kendall correlation with the *corr.test* function for each soil compartment. Then combinations giving significant correlation (*p. value* < 0.05) were further analysed using generalised additive mixed models with a Gaussian family and identity link, as yielding the best residual diagnostics, and best models were selected using AIC.

The effect of the health status on mean enzymatic activities was tested using a generalised additive mixed model for each enzymatic activity as response variable, and soil type, health status and their interaction as fixed effects with a Gaussian family and identity link. Tree pair identity was considered as a random factor in the models. Pairwise comparison between soil types and health status were performed and *p* values adjusted as previously described. To specifically test for links between laccase activity and the relative abundance of *Russula* as a function of health status, we performed a generalised additive mixed model considering laccase activity as response variable and rarefied abundance of *Russula* and health status and their interaction as fixed effects with a Gaussian family and identity link. Tree pair identity was considered as a random factor in the model. Kendall correlation between rarefied abundance of *Russula* in non-declining oak rhizospheric samples and oak declining rhizospheric samples and laccase activity were calculated using the *stat_cor* function of ggplot.

### Data availability

Raw data were deposited in the NCBI Sequence Read Archive (SRA) under SRA accessions numbers SAMN56358317 to SAMN56358490 for the 16S bacterial data, SAMN56432516 to SAMN56432689 for the ITS fungal data and SAMN56433258 to SAMN56433431 for the oomycete data (project PRJNA 1419566). All R scripts are available on demand.

## RESULTS

### Fungal and bacterial communities of non-declining trees showed no major signs of dysbiosis

Fungal and bacterial communities of the non-declining trees were mainly structured by compartment (BS, Rh or R) and by the plot soil type (results of the PERMANOVA, **Table 1A**, **Suppl. Figure 1A, B**). Fungal richness was reduced by half in roots compared to the bulk soil and by 15% in rhizospheric samples no matter the soil type (Poisson GAMM, *p* value < 0.01, **Suppl. Figure 1C**). Conversely, bacterial richness slightly varied depending on soil type, compartment and tree circumference (Poisson GAMM, *p* value < 0.05). While both communities were dominated by a few dozen genera in the three compartments, the relative abundance of certain taxa varied greatly according to soil type (**Figure 2A, B**).

**Figure 2.**
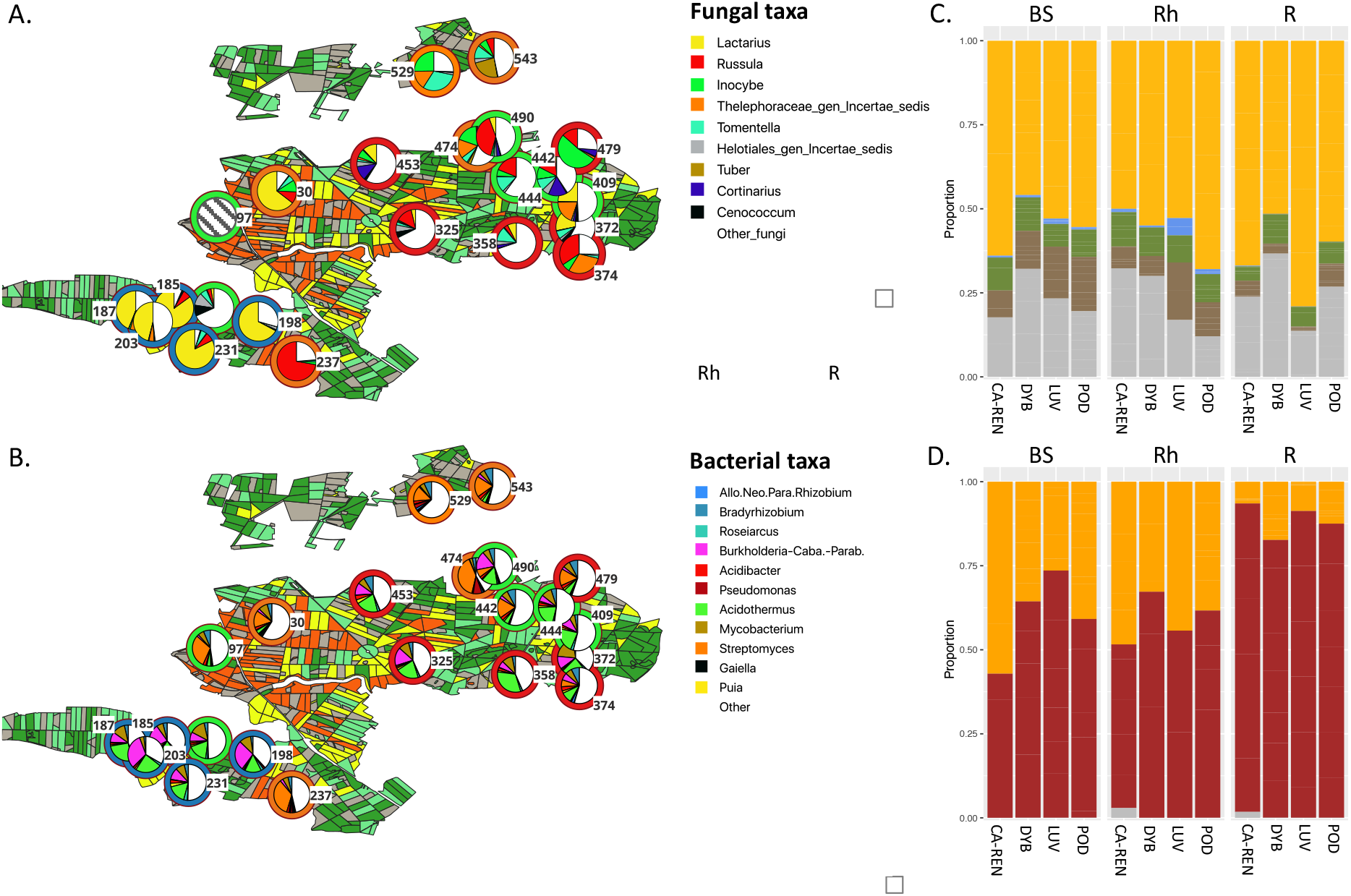
Distribution and relative abundances of the dominant genera of fungi (A) and bacteria (B) detected in root samples of non declining oaks. Color of the outer circle of the pie chart indicate the soil type: : orange-DYB, red – CA-REN , blue – LUV, green – POD. Dieback level, as determined through DEPERIS method, is depicted for the entire Chantilly forest from no mortality (green) to high mortality (red). Numbers indicate sampled plots. Grey hatched pattern : no data. **C.** Proportion of fungal trophic guilds as a function of soil types and compartments. Orange: ECMs, blue: plant pathogens, green: saprotrophs, brown: others, grey: unknown. **D**. Proportion of high (orange) and low biomass (red) ECM exploration types as a function of soil types and compartments.

**Table 1.**
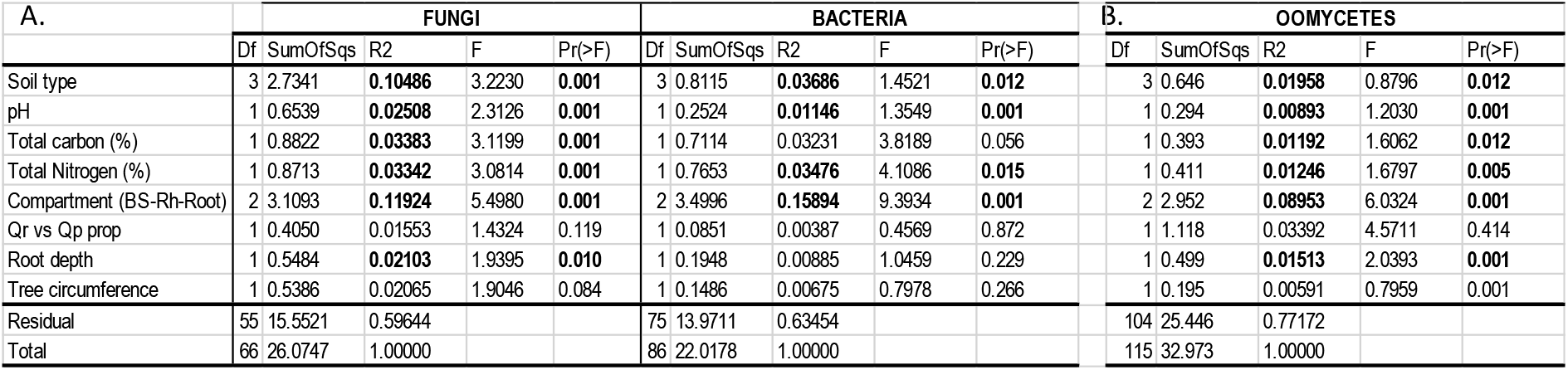
PERMANOVA based on Bray-Curtis (A. fungi, bacteria) or Jaccard (B. Oomycetes) dissimilarity matrices showing principal abiotic and biotic drivers of fungal, bacterial community and oomycetes structures of non diseased trees. Plots were considered as random effects. Significant values (*p* < 0.05) are highlighted in bold. Only samples in which Oomycetes were detected were considered for structure analyses (113 out of 174 samples). Qr vs Qp prop: Proportion of *Q. robur* vs *Q. petrea* in the plots based on genetic analyses (Sénéchal-Chevalier et al. 2024)

| A. |  | **FUNGI** |  |  |  |  | **BACTERIA** |  |  |  |  | B. **OOMYCETES** |  |  |  |
|---|---|---|---|---|---|---|---|---|---|---|---|---|---|---|---|
|  | Df | SumOfSqs | R2 | F | Pr(>F) | Df | SumOfSqs | R2 | F | Pr(>F) | Df | SumOfSqs | R2 | F | Pr(>F) |
| Soil type | 3 | 2.7341 | **0.10486** | 3.2230 | **0.001** | 3 | 0.8115 | **0.03686** | 1.4521 | **0.012** | 3 | 0.646 | **0.01958** | 0.8796 | **0.012** |
| pH | 1 | 0.6539 | **0.02508** | 2.3126 | **0.001** | 1 | 0.2524 | **0.01146** | 1.3549 | **0.001** | 1 | 0.294 | **0.00893** | 1.2030 | **0.001** |
| Total carbon (%) | 1 | 0.8822 | **0.03383** | 3.1199 | **0.001** | 1 | 0.7114 | 0.03231 | 3.8189 | 0.056 | 1 | 0.393 | **0.01192** | 1.6062 | **0.012** |
| Total Nitrogen (%) | 1 | 0.8713 | **0.03342** | 3.0814 | **0.001** | 1 | 0.7653 | **0.03476** | 4.1086 | **0.015** | 1 | 0.411 | **0.01246** | 1.6797 | **0.005** |
| Compartment (BS-Rh-Root) | 2 | 3.1093 | **0.11924** | 5.4980 | **0.001** | 2 | 3.4996 | **0.15894** | 9.3934 | **0.001** | 2 | 2.952 | **0.08953** | 6.0324 | **0.001** |
| Qr vs Qp prop | 1 | 0.4050 | 0.01553 | 1.4324 | 0.119 | 1 | 0.0851 | 0.00387 | 0.4569 | 0.872 | 1 | 1.118 | 0.03392 | 4.5711 | 0.414 |
| Root depth | 1 | 0.5484 | **0.02103** | 1.9395 | **0.010** | 1 | 0.1948 | 0.00885 | 1.0459 | 0.229 | 1 | 0.499 | **0.01513** | 2.0393 | **0.001** |
| Tree circumference | 1 | 0.5386 | 0.02065 | 1.9046 | 0.084 | 1 | 0.1486 | 0.00675 | 0.7978 | 0.266 | 1 | 0.195 | 0.00591 | 0.7959 | 0.001 |
| Residual | 55 | 15.5521 | 0.59644 |  |  | 75 | 13.9711 | 0.63454 |  |  | 104 | 25.446 | 0.77172 |  |  |
| Total | 66 | 26.0747 | 1.00000 |  |  | 86 | 22.0178 | 1.00000 |  |  | 115 | 32.973 | 1.00000 |  |  |

The fungal community was dominated by ECM fungi (∼60% in average) followed by saprophytes and plant pathogens (**Figure 2C**). ECM communities were largely dominated by “low biomass” exploration types in the roots, representing 87.9% of the ECM (+/-2.4%) regardless of soil type (**Figure 2D**). The distribution of trophic guilds was similar in all soil types except for plant pathogens. The latter were more abundant in the rhizosphere in LUV soils, reaching 5% of the reads, while they represented about 1% in other soils. Nevertheless, these taxa were primarily leaf pathogens, not root pathogens (**Suppl. Table 3**). None was detected in the root samples.

Eight enzymatic activities involved in organic matter degradation were measured in the rhizospheric soil of non-declining trees. Positive but highly variable signals were obtained for seven of them: β-glucosidase, cellobiohydrolase, xylosidase, glucuronidase, N-acetylglucosaminidase, acid phosphatase and laccase (**Suppl. Figure 2**). Only cellobiohydrolase activity differed between soil types. This activity was on average higher in LUV than in CA-REN soil (**Suppl. Figure 2**, Gaussian GAMM, *p* value = 0.05). *Quercus* species, together with nitrogen percentage and *Lactarius* relative abundance explained 75% of the variance of cellobiohydrolase activity (*p* value < 0.001, **Suppl. Table 4**). This activity was higher in the rhizospheric soil of *Q. petrea* than *Q. robur* and was positively correlated with the relative abundance of *Lactarius* (*p* value < 0.05, Kendall correlation test). None of these variables were correlated with each other (data not shown). The same method was applied to investigate relationships between enzymatic activities and the relative abundance of the 12 most dominant ECM fungi in the rhizosphere. Laccase activity was strongly determined by the proportion of *Russula* and of fine roots (Gaussian GAMM, R^2^ = 0.68, *p* value < 0.01, **Suppl. Table 4**). Laccase activity was positively correlated with the proportion of *Russula*, while negatively correlated with the proportion of fine roots in soil. Finally, glucuronidase activity could not be associated with ECM species but was partially determined by *Quercus* species (Gaussian GAMM, R^2^ = 0.48, *p* value < 0.001, **Suppl. Table 4**) and was higher in the rhizospheric soil of *Q. petrea* than *Q. robur*. No relationship was found for other enzymatic activities nor between dominant non-ECM fungi and enzymatic activities (data not shown).

### Oomycetes were frequently detected in samples from non-declining trees

ASVs assigned to oomycete species could be detected in 58 (i.e., 66%) of the 87 samples associated with non-declining trees, after bioinformatic treatments. There was no difference in the number of oomycete positive samples between compartments (**Suppl. Table 5**, Khi² test, *p* value = 0.09). By contrast, it differed between soil types: more samples were positive in CA-REN and POD soils than in DYB and LUV within the same soil compartment (**Suppl. Table 5**, Khi² test, *p.* value = 0.05 for R, Rh and BS).

The structure of the oomycete communities significantlydiffered between compartments (PERMANOVA, R² = 13.8%, *p* value < 0.001, **Table 1B**). The other tested parameters had no significant effect on the composition of oomycete communities associated with non-declining trees. The most frequent *Phytophthora* species associated with non-declining trees was *P. quercina* (detected in 66, 52 and 79 % from the BS, Rh and R samples, respectively, **Figure 3**) followed by *P. psychrophila* (in 48%, 45% and 55% from the BS, Rh and R samples, respectively) and the multi-affiliation taxon *P. borealis-riparia* (in 45%, 45% from the BS and the Rh of the ND trees, respectively and 55% for the R) and *P. borealis-riparia-fragaefolia* (in 31%, 17% and 45% from the BS, Rh and R of the ND trees, respectively).

**Figure 3.**
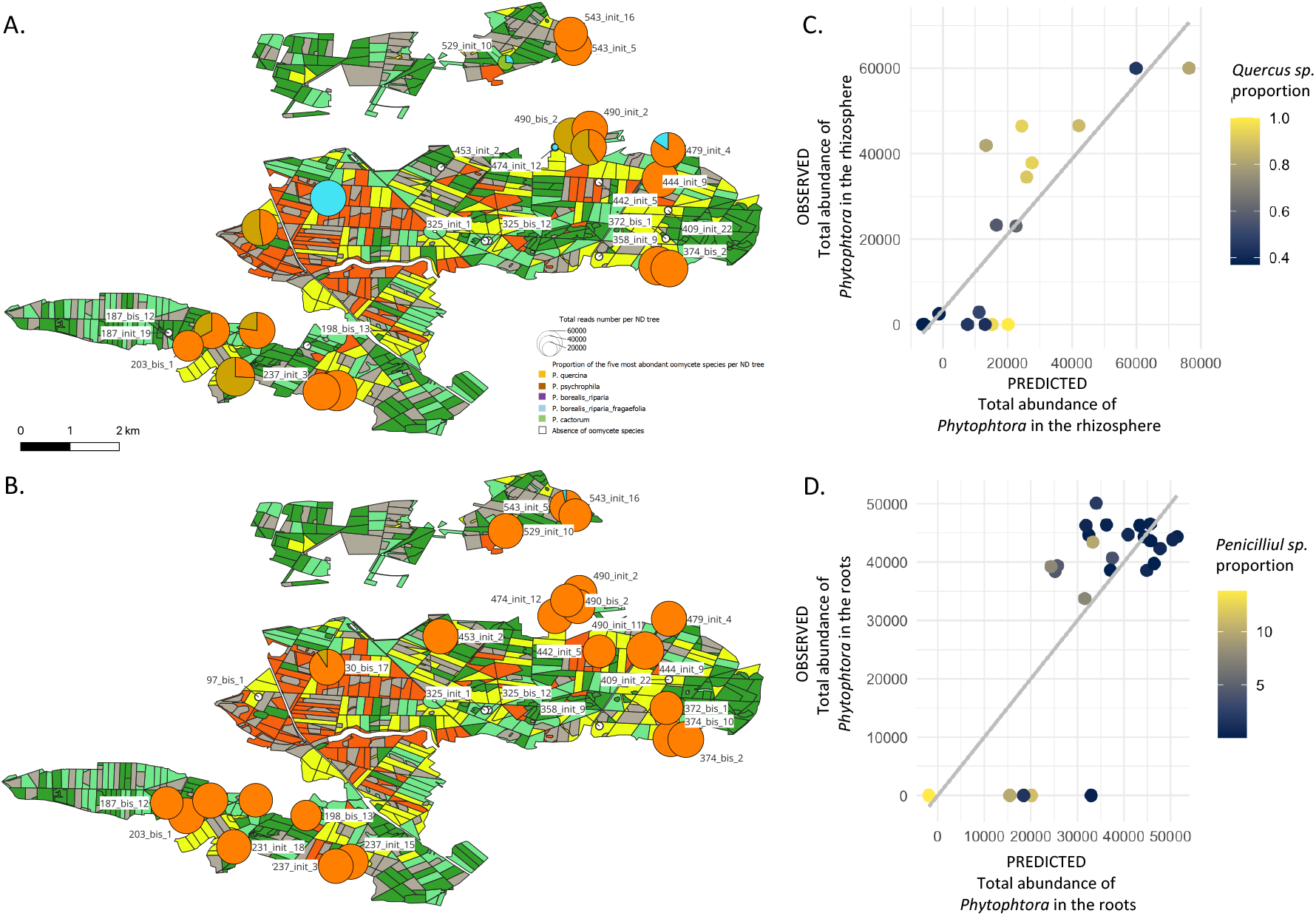
Distribution of oomycestes across space and compartments. **A, B.** Maps of the proportion of the abundance of the main oomycetes species in ND oaks in bulk soil (A) and in roots (B) compartment. **C. D.** Scatter plots comparing predictions of GAMM models of total abundance of *Phytophtora* in the rhizosphere (C) and the roots (D) of non declining oaks based, using plots as random effects. C. Fixed effects: *Bacillus* proportion*, Quercus* proportion*. D*. Fixed effects: *Penicillium* and *Acidothermus* proportions.

Oomycetes species were detected with the same frequency both in *Q. robur* and *Q. petraea* samples, whatever the compartment (Khi² test, *p* value = 0.87). We further tested whether *Phytophthora* abundance can be influenced by the presence of the most abundant other microorganisms, using Spearman correlation tests (**Suppl. Figure 3**). No correlation was found in the bulk soil compartment (data not shown) but both positive and negative correlations were found between the total number of reads of *Phytophthora* and several fungal and bacterial taxa in the rhizosphere and roots, respectively. We further explored the relationship between microbial taxa, taking into account the available environmental, tree and stand variables and we found that the total number of reads of *Phytophthora* in the rhizosphere of oak roots can be largely explained by the number of reads of the bacterial taxa *Bacillus* and the proportion of oak trees in the stand (R^2^ = 0.54). The model predicts higher levels of *Phytophthora* colonisation of the rhizosphere oak trees when the relative abundance of *Bacillus* in the rhizosphere is above 10% and the proportion of oaks in the stand is high (**Figure 3C**). Conversely, the total number of reads of *Phytophthora* in the roots of oaks was negatively correlated with the abundance of the fungus *Penicillium* (R^2^ = -0.55, *p* value < 0.01) and, to a lesser extent, the bacterium *Acidothermus* (R^2^ = -0.44, *p* value < 0.01). A GAMM model combining *Penicillium* and *Acidothermus* abundances in the roots with tree circumference explained 46% of the variance in *Phytophthora* abundance in oak roots, although the significance of tree circumference was weak (*p* value = 0.07, **Figure 3D**). None of the environmental variables improved the models.

### Bacterial and fungal communities reacted differently to oak decline

Since compartment (BS, Rh, R) and soil types were strong drivers of microbial communities’ composition in non-declining trees, analyses were performed for each sample compartment and on each soil type separately in order to search a possible compartment-associated response to oak decline. Since sequencing depth was insufficient for some fungal samples, 26 pairs of samples were discarded for the analysis of fungal communities and soil types. Only soil types for which at least three pairs of samples remained were considered. Consequently, fungal BS and Rh samples taken on CA-REN and LUV soils could not be analysed (see **Suppl. Table 2**).

Bacterial and fungal communities reacted differently to the health status of the oak trees. The tree health status had no significant influence on the fungal community structure independently of the compartment and the soil type (Bray-Curtis matrix, PERMANOVA test – **Suppl Table 6**). By contrast, a small but significant effect of the health status was detected for the bacterial communities in the rhizosphere (R^2^ = 1.1%, *p* value = 0.05) and the bulk soil (R^2^ = 4.5%, *p* value = 0.04, **Suppl. Table 6**). Response was more pronounced on POD soils as illustrated by constrained analysis of principal components (**Figure 4A**). POD-associated bacterial communities were characterized by higher richness in response to decline in BS and a lower richness in the Rh and the R of declining oaks (Poisson GAMM, *p* < 0.01, **Figure 4B**).

**Figure 4.**
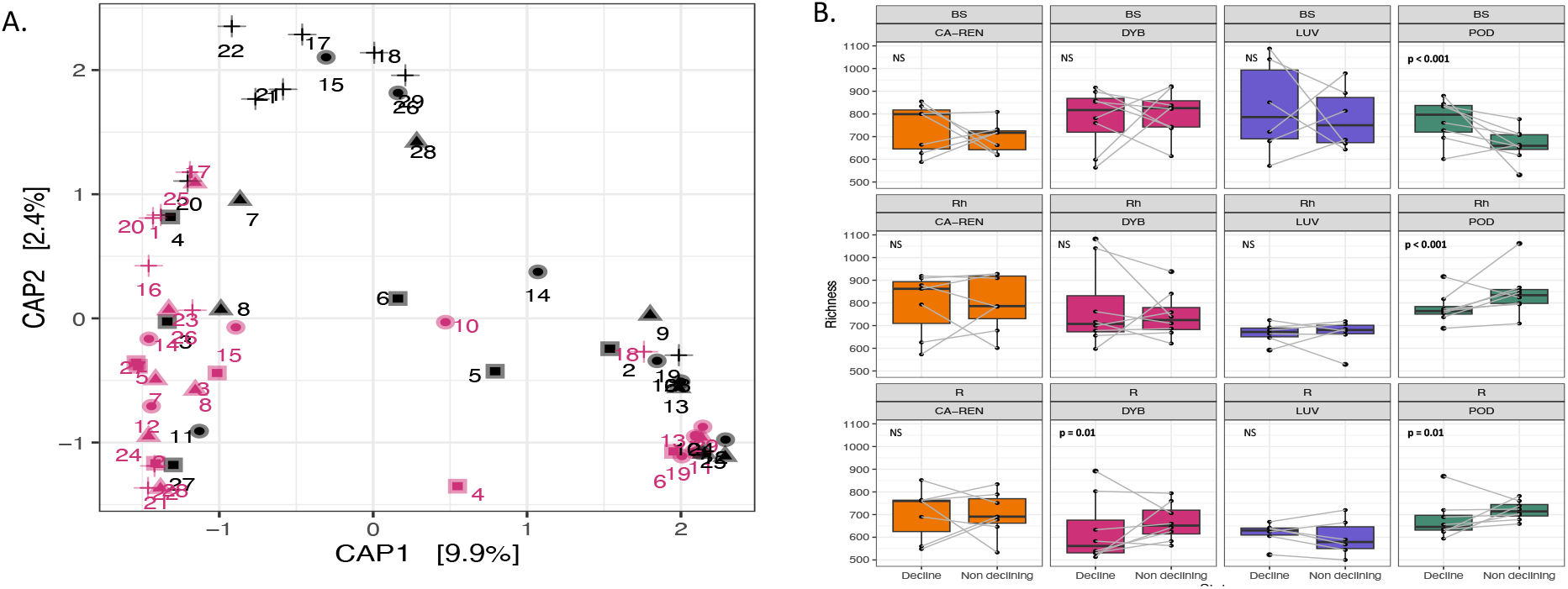
Relationships between oak health status and bacterial communities. CAP ordination using Bray-Curtis dissimilarity matrix using health tree status and soil type as constrains. Numbers correspond to each pair of samples. Shapes indicate the soil type: DYB – circle, CA-REN – triangle, LUV – square, POD – star, colors indicate health status: red – non diseased, black: diseased. **B.** Richness as defined by total number bacterial ASVs. Paired trees are connected by grey lines. Generalized mixed model using health status x soil type as fixed effects and pair as random effects. Poisson distribution was used to test the combined effects soil type and health status on richness. BH method was used to correct p values for multiple testing. Corrected *p.* values < 0.01 were considered significant.

Regarding fungi, the overall comparison at the guild level showed no differences in the relative abundance of fungal guilds between ND and D trees, regardless of the soil type or compartment. Furthermore, the proportion of plant pathogens was low and around 1% and was similar in ND and D samples, regardless of sample type or soil type (**Figure 5A**). However, the proportion of ECM high biomass exploration types was reduced by half in the root samples of declining trees, dropping from 11.8 +/- 2.5% to 6.0+/-1.3% (*p* value < 0.01 Beta GAMM, **Figure 5B**). Fourteen ECM genera recorded as “high biomass” were detected across the R samples. Among those, four showed significant variation of relative abundance between roots of declining and non-declining oaks: *Amanita, Piloderma, Xerocomellus* and *Melanogaster* (**Figure 5C**, *p* value < 0.01, negative binomial GAMM model). While the relative abundance of the three first ones decreased in the roots of declining oaks, the relative abundance of *Melanogaster* increased.

**Figure 5.**
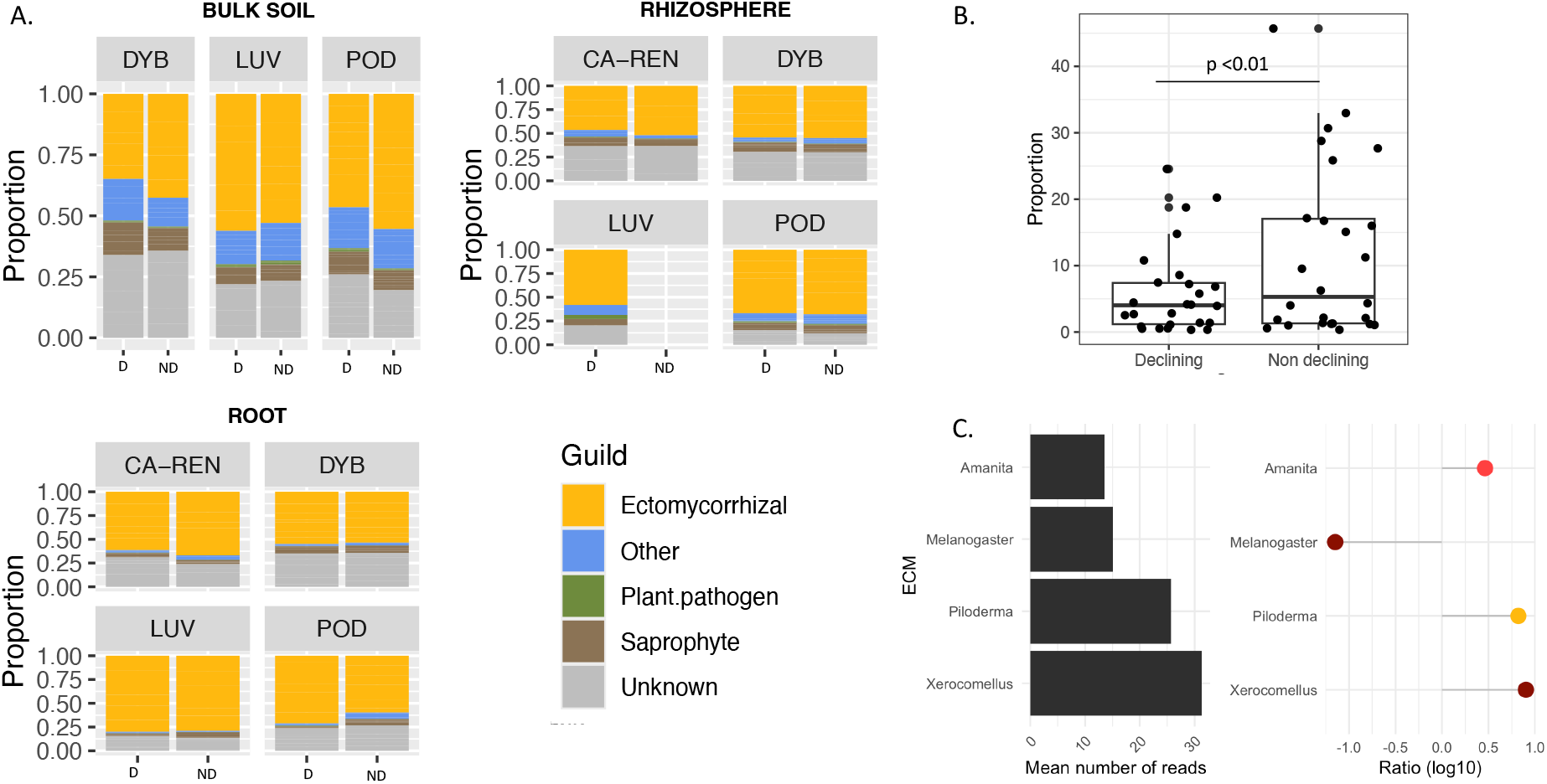
**A**. Average proportions of main fungal trophic guilds of non-declining vs declining oaks as a function of soil type. Not enough replicates passed quality control to be analysed for bulk soil CA-REN and LUV-ND rhizosphere samples **B.** Proportion of high biomass ECM types as a function of tree health status. Generalized mixed model using sanitary status x soil type as fixed effects and pair as random effects with a Beta distribution was used to test the combined effects soil type and status on high biomass proportion. **C**. Mean number of reads and log fold change of relative abundance between non decline and decline status for fungal taxa of high biomass exploratory type significantly enriched in samples of non declining or declining trees. Generalized mixed model using sanitary status + soil type as fixed effects and pair as random effects with a non binomial distribution. Colors indicate taxonomical order. brown: Boletales, yellow: Atheliales, red: Agaricales.

### Oomycete communities differ between declining and non-declining oaks

The tree health status and the plot soil type had a significant effect on the beta diversity of the oomycete communities but only in the root compartment, the health status explaining 7% of the distance between communities and the plots 11% (*p* value < 0.01, **Suppl. Table 6**). Oomycetes were present at a higher rate in R and BS samples (80% positive samples) than in the Rh of D tree samples (45%, **Suppl. Table 7**). *P. quercina* was the most frequent and abundant oomycete identified but it was detected in similar proportions in both ND and D trees (23 pairs for the BS and R compartments, 13 for the Rh). By contrast, oak health status was correlated with the number of reads of *Phytophthora plurivora* (z = 2.56, *p* = 0.011; **Figure 6A**) and marginally with the number of reads of *P. cactorum* (z = 1.76, *p* = 0.078). However, their detection was rare (i.e. 10 to 30% of samples, **Suppl. Table 7**).

**Figure 6.**
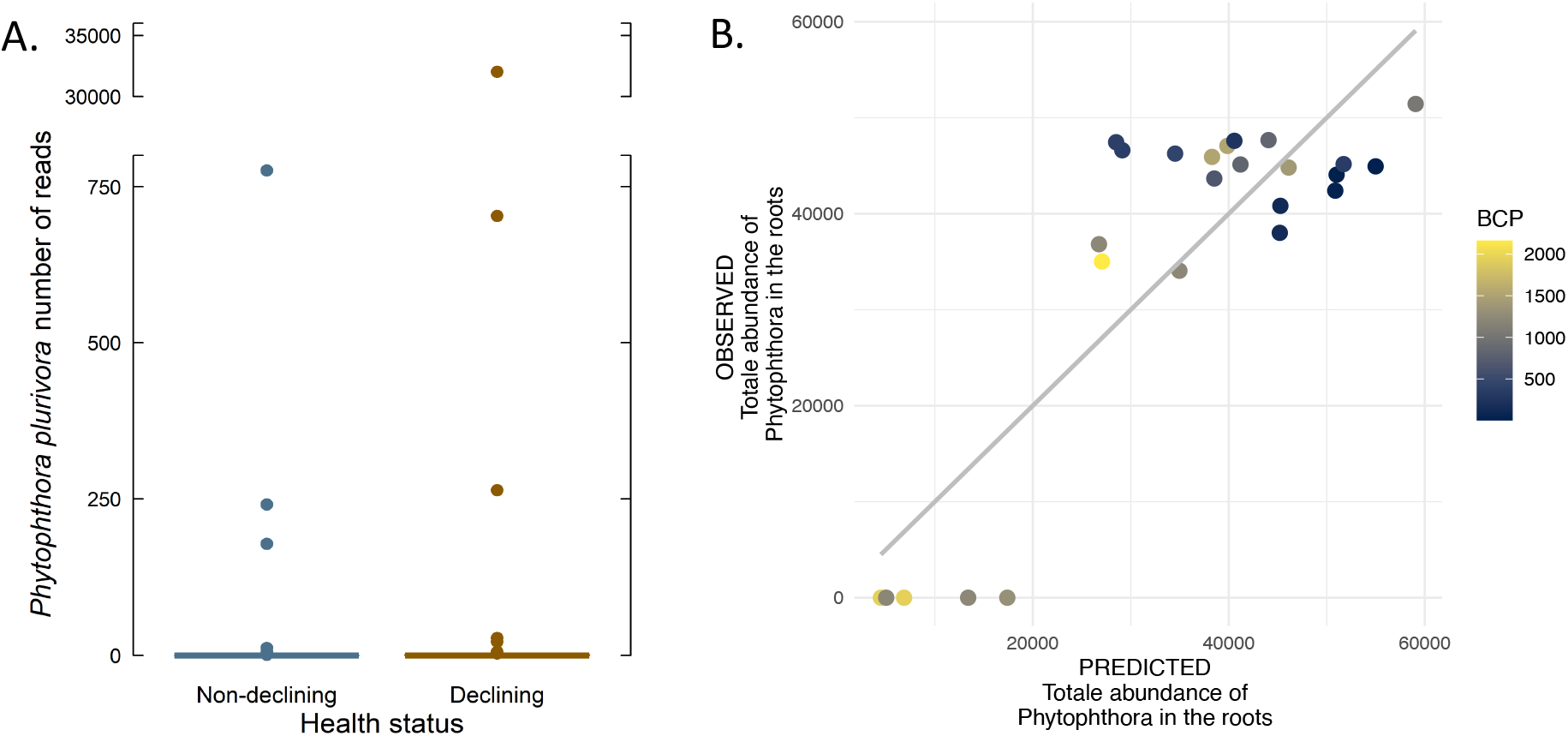
Distribution of oomycestes as a function of health status. **A.** Number of reads of *Phytophthora plurivora* as a function of the health status the roots. Optimal binomial GAMM model using plots and pairs as random effects. **B.** Scatter plot comparing prediction of gaussian GAMM model of total abundance of *Phytophtora* in declining oak roots as a function of the relative abundance of the *Burkholderia-Caballeronia-Paraburkholderia* taxa (BCP), the percentage of N in soil, tree circumference, the proportion of oak in the plots and the species of oak, using plots & pair as random effects.

Interactions between microorganisms, and notably *Phytophthora,* may be modified in the roots of declining trees compared to non-declining trees. The abundance of *P. quercina* but not *P. plurivora* and *P. cactorum,* was positively correlated with the abundance of *Conexibacter* (R^2^= 0.51, *p* value < 0.01) and *Acidothermus* bacteria (R^2^ = 0.45, *p* value < 0.01) in the bulk soil of declining oaks. A weak positive correlation was found between the abundance of *Russula* and of *Phytophthora* in the rhizosphere of declining oaks (R^2^ =0.32, *p* value < 0.01). A GAMM model combining *Russula* abundance in the rhizosphere with the percentage of N in soil explained 36% of the variance in *Phytophthora* abundance in the oak rhizosphere, although the significance of N level was marginal (*p*. value = 0.06). By contrast, the variability of *Phytophthora* abundance in oak roots could be rather well explained (R^2^ = 0.66) by the relative abundance of the *Burkholderia-Caballeronia-Paraburkholderia* taxa (BCP), the percentage of N in soil, tree circumference, the proportion of oak in the plots and the species of oak (**Figure 6B**). The proportion of BCP in the roots and the proportion of oaks in the plot had a negative effect on the abundance of *Phytophthora*, while the other parameters had a positive effect.

### Microbial activities were partially affected by tree health status

We further tested whether the functioning of microbial communities, in particular ECM communities, was altered by the health status of the oak trees through the measurement of enzymatic activities in the rhizospheric soil samples. Effects of the health status in interaction with the soil type were only found for two enzymatic activities: the N-acetylglucosaminidase activity which is involved in the degradation of fungal and insect chitin and the laccase activity which is involved in lignin degradation from plant cell walls. Both activities were reduced in the rhizospheric soils of declining trees compared to healthy trees, but the effect was soil dependent and was only detectable in LUV soil for chitinase activity and in DYB soil for laccase activity (**Figure 7A, B**). Since laccase activity correlated with the relative abundance of *Russula* in non-declining trees (**Suppl. Table 4**), we looked for a possible interaction between the health status and the relative abundance of this ECM. Despite being still detected in the rhizosphere of declining oaks, the relative abundance of *Russula* was no longer correlated with laccase activity (**Figure 7C**). Instead, little to no laccase activity was measured in the rhizosphere of declining oaks despite variable levels of colonization by *Russula*, suggesting a potential reduction of the activity of the ECM when oak trees are declining. No correlation was found with the relative abundance of *Phytophthora* (data not shown).

**Figure 7.**
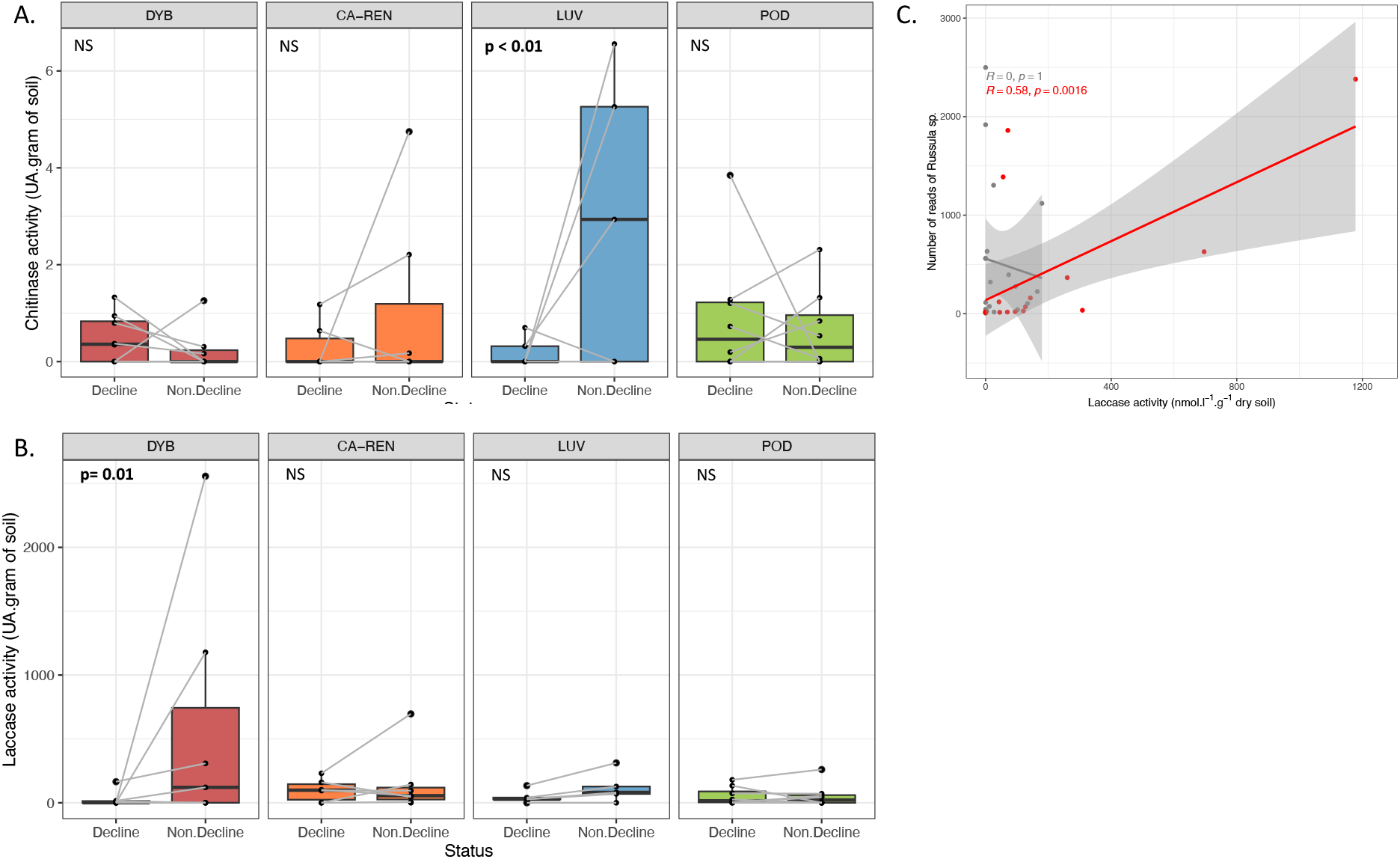
A. N-acetylglucosaminidase and B. Laccase activities in rhizospheric soil of non-declining and declining oaks as a function of soil types. Generalized mixed model using sanitary status x soil type as fixed effects and pair as random effects with a Gaussian distribution was used to test the combined effects soil type and status on enzymatic activities. BH method was used to correct p values after post-hoc comparisons. Corrected p.values ≤ 0.01 were considered significant. Lines indicate paired values. C. Relationship between laccase activity and *Russula* sp. abundance in the rhizospheric soil of declining (grey) and non declining oaks (red). Linear regressions using Kendall correlations are given for both heatlh status: red = non-declining, grey = declining.

### Integration of soil edaphic properties, tree parameters and microbial communities to predict tree health status

In order to obtain an integrated vision of the variables which best explained the health status of the oak trees, we used Elastic Net approach. Among the 6 365 parameters tested, 22 variables identified by Elastic net were validated by binomial GAMM and all were bacterial ASV (**Suppl. Table 8**). Eighteen corresponded to the relative abundances of bacterial ASV in BS samples, while only four were associated with Rh and R (**Figure 8A**). Proteobacteria and Actinobacteria dominated, ASV from the latter being mainly enriched in samples from declining trees. Conversely, explanatory bacterial ASV of the non-declining status belonged essentially to Proteobacteria.

**Figure 8.**
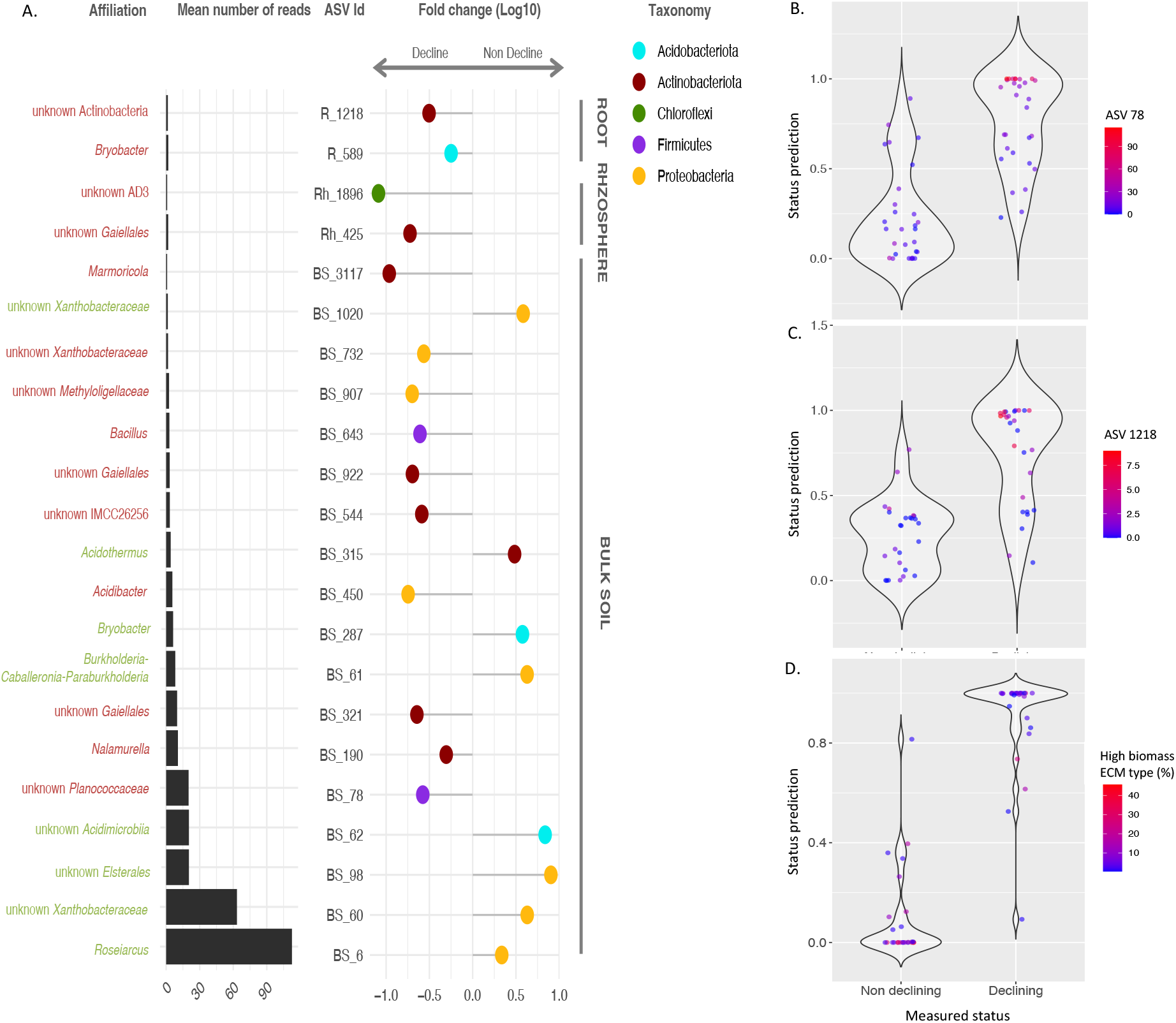
Integration of microbial and non microbial variables to predict health status. **A.** Mean number of reads and log fold change of the relative abundance of bacterial taxa between non declining and declining trees enriched in samples according to Elastic net analysis followed by GAMM validation (corrected *p.* value <0.01). **B.** Prediction of tree health status based on generalised mixed model predicting status as a function of the relative abundance of ASV78 and 62 in bulk soil of the trees. (p<0.01, R2 = 0.48). **C.** Prediction of tree health status based on generalised mixed model predicting status as a function of the relative abundance of ASV880, 1218 and the proportion of high biomass ECM exploratory types in the root samples (p<0.01, R2 = 0.45). **D.** Prediction of tree health status based on generalised mixed model predicting status as a function of the relative abundances of ASV 62 and 78 in bulk soil combined with ASV 1218 and the proportion of high biomass ECM exploratory types in the root samples (p<0.01, R2= 0.79). Each dot represents the predicted value health status (0 = non declingin, 1=declining) of samples known to be non declining or declining (X-axis). Gradual colors indicate the relative abundance of ASV or high biomass ECM measured in the samples from low (blue) to high (red).

To go further, as the health status of oak trees was found to correlate with changes in the relative abundance of several microbial taxa, microbial community parameters (e.g., bacterial alpha diversity and ECM functional exploration types) and enzymatic activities as a function of soil properties, we tested whether the health status of trees could be modelled using these variables with a GAMM model. Three models were established, one per compartment (BS, Rh and R). Soil edaphic properties and available plot tree parameters, neither the abundance nor the presence of oomycete taxa, when considered individually or collectively, were able to predict tree health status (data not shown), in contrast to other microbial variables (**Suppl. Table 9**). Best models were obtained with BS and R data (R^2^= 0.48, Figure 8B and 0.45, **Figure 7C**) but combination of BS (ASV 62 & 78) and R (ASV 1218 & high biomass exploration type) variables strongly improved predictions (R^2^ =0.79, **Figure 8D**).

## DISCUSSION

Soil and tree associated microbiota can have a dual effect: on the one hand, they can protect trees against stressors; on the other hand, they can amplify the effect of stressors through pathogenic activities and/or a reduction of their beneficial activities. Conversely, as trees are an important source of carbon and metabolites for soil- and root-associated microbiota, tree decline may negatively affect the microbiota and lead to negative feedback loops, which can further amplify the unhealthiness of the ecosystem (Manion 1981, Albornoz et al. 2025). In this study, we aimed at testing whether altered soil microbiota could be a contributing factor for decline and whether chronic oak decline affects soil and root microbiota composition and activities for nutrient cycling. We hypothesized that these effects would be different depending on soil types as microbial assemblages associated with oaks display significant spatial variation across forest sites (Suz et al. 2014, Labouyrie et al. 2022).

### Soil and oak associated microbiota showed no sign of dysbiosis in non-declining trees

In a first step, we explored whether the microbial communities associated with the oak trees that were considered healthy at the time of sampling showed any abnormalities that could indicate microbial predisposing factors for chronic oak decline in the Chantilly forest. We detected no signs of dysbiosis in the root and soil microbiota at the level of fungal and bacterial communities sampled from healthy trees, neighbours of declining trees and characterized with global ITS and 16S metabarcoding markers. Indeed, the compositions and functions of these communities were very similar to those described for European healthy oak forests (Suz et al. 2014, Labourie et al. 2022) and responded to the expected drivers in terms of compartment effect, tree species and physico-chemical properties (Lladóet al. 2018). The fungal communities were largely dominated by ECM, while the level of pathogens was very low. No primary or opportunistic fungal pathogens of oak were detected, such as *Armillaria* or *Collybia*, or bacterial species associated with acute oak decline (Denman et al., 2018). However, 12 species of *Phytophthora* in both soil and roots of non-diseased trees were detected. Among those, several ones are considered as potential oak pathogens including *P. quercina* (Jung et al. 1999b), *P. cactorum* (Jonsson 2004), *P. psychrophila* (Jung et al. 2002), *P. pseudosyringae* (Jung et al. 2003) and *P. plurivora* (Jung & Burgess 2009), which can cause fine root rot in oak seedlings. Nonetheless, these results are in line with other surveys of *Phytophthora* communities in European oak forests which all highlight the ubiquity and diversity of *Phytophthora* communities (Vettraino et al. 2002, Riddell et al. 2019, Hansen and Delatour 1999, Jankoviak et al. 2014, Lopez Garcia et al. 2025). The most frequently identified oomycete species in our samples taken from non-declining trees was *P. quercina*, regardless of the compartment analysed. We found it to be ubiquitous, with its abundance, assessed by the number of reads, unrelated either to the tree health status (p=0.41) or to the proportion of fine roots (data not shown). Thus we advocate that the presence of *P. quercina* is likely not a sign of dysbiosis. However, since *P. quercina* has been associated to greater fine root damage and an increased risk of developing severe crown symptoms (Jung et al. 2000), we can not rule out the hypothesis that non-declining trees, colonized by *P. quercina*, may be at higher risk of developing severe decline, as suggested by López Garcia et al. (2025).

### Oak decline had a differential effect on root and bulk soil microbiota

Overall, chronic oak decline had little effect on microbiota at the community level in the Chantilly forest. No difference could be detected between fungal communities of declining vs non declining trees, no matter the compartment or the soil type considered. A slight but significant effect was measured for bacterial communities, particularly on POD soil, but it was not consistent across compartments: in bulk soils, there was a shift towards richer bacterial communities in declining oak trees, but this shift was reversed in oak roots and rhizosphere. These results contrast with those of López-Garcia et al. (2025), who found that fungal, but not bacterial, communities were affected by chronic oak decline. However, our results are in line with studies on the relative response of fungi and bacteria communities to drought stress: fungal communities showing most often a greater tolerance to water deficit than bacterial ones (Bardgett et al. 2020, de Vries et al. 2020). The discrepancies with López-Garcia et al. (2025) may come from soil type since most effects we observed were found on POD soil, but the López-Garcia report lacks soil characteristics. In addition, some authors have suggested that old forests, such as the Chantilly forest, where declining and vigorous trees grow close together, may keep an abundant resource of ECM fungi thanks to the healthy trees which would provide relative inertia in response to disturbance (Kovacs et al. 2020). Such phenomenon may explain why we found little differences between the fungal communities of declining vs non declining trees in the present study.

Nevertheless, the fact that an opposite response was found between bulk soil vs rhizospheric and root samples could suggest that chronic oak decline induces differential direct and indirect effects on the soil microbiota. Bulk soil is not, or far less, under the influence of the tree physiology as it does not receive root exudates and is thus largely enriched in oligotrophic bacteria (Uroz et al. 2016). Bacterial communities in this compartment are thus more prone to be directly affected by environmental variations. Crown reduction resulting from tree decline induces modification of soil edaphic conditions by reducing leaf litter on one side and increasing direct exposure to heat and drought on the other side (Anderegg et al. 2012, Garcia-Angulo et al. 2020). Such stresses imposed on various European soil microbiomes produced a consistent response across soils which was phylogenetically conserved: fungi as well as Acidobacteria, Firmicutes, Actinobacteria and Bacteroidetes were in majority resistant to these stresses while Proteobacteria were rather sensitive (Knight et al. 2024). In line with these results, a large part of the taxa enriched in the BS samples of the declining trees belonged to the Firmicutes and Actinobacteria phyla, while those associated with the non-declining trees, which strongly negatively responded to the decline, mainly belonged Proteobacteria. Our results therefore suggest that tree decline may indirectly lead to increased drought and heat stress in soils by opening the canopy.

Chronic oak decline also partly impacted rhizospheric and root-associated taxa. Although the vast majority of root colonising taxa did not seem to be affected by the health status of oaks, the 50% drop in high-biomass ECM exploration types linked to declining oak roots, along with reduced bacterial alpha diversity, may reflect altered root–microbiota interactions caused by changes in tree physiology. In previous studies on cases of severe oak decline caused by pathogen infections, the rapid and massive loss of vitality was associated with an alteration of root microbiota, including ECM, suggesting a direct connection between microbiota composition and tree physiology (Pinho et al 2020, Gomez Aparicio et al 2022). By contrast, chronic oak decline was previously reported to have moderate effects on mycorrhizal colonization rates, sometimes resulting in species replacements, suggesting, in this cases, more limited effects on the root-microbiota interactions than for AOD (Causin et al. 1996, Kovacs et al. 2000, Garcia Angulin et al. 2020, Lopez Garcia et al. 2025). Flux and carbon allocation studies are needed to understand how oak decline affects the root microbiota.

High biomass ECM fungi are characterised by their medium-to long-distance exploration types. They are known as “high biomass” because they establish extensive mycorrhizal networks in the soil, and they are considered to require more plant carbon for mycelial growth (Hobbie & Agerer 2010, Ekblad et al. 2013, Lilleskov et al. 2019). Whether declining oaks associated less with C-expensive ECM or high biomass ECMs colonized less declining oaks remains to be determined. Nevertheless, this potentially likely impacts mineral and water nutrition of the declining trees as high biomass ECMs tend to explore and extract nutrients from deeper soil horizons (Hobbie et al. 2013, Khalfallah et al. 2024).

### Roots of declining oaks presented higher colonisation by two species of Phytophthora compared to healthy trees

Contrary to our initial hypothesis, chronic oak decline was not associated with an increased colonisation by oak root pathogens such as *Armillaria* or *Collybia*. However, despite their rare occurrence, the oak pathogens *P. cactorum* and *P. plurivora* were positively associated with the declining status of the trees. Jankoviac et al (2014) previously reported that *P. plurivora* was more frequently isolated from declining oak trees in Polish forests than visibly healthy trees. Whether these *Phytophthora* species directly contribute to oak decline and/or can more easily colonize roots of declining oaks remains to be determined. Most studies addressing this question are performed on oak seedlings and can be hardly extrapolated to adult trees in natural ecosystems, where they can interact and assembly with other microbial species (Jung et al. 1999, Jönsson 2004, Jankoviak et al. 2014).

Our study provides clues on the combination of biotic and abiotic factors which may favour differently the development of *Phytophthora* species in asymptomatic and declining oak trees. The combination of four factors were at play in our data: microbial communities, soil properties, tree parameters and forest management. None of these factors independently predicted *Phytophthora* colonisation, but their combination did. Declining trees were more likely to be colonized when they were aged *Q. robur* growing on a soil rich in nitrogen, while the presence of members of the *Burkholderia-Cabaleronia-Paraburkholderia* in abundance was detrimental to *Phytophthora* development in the roots. A higher proportion of oaks in the stand was also associated with a reduced presence of *Phytophthora*, which suggests that species mixing, as opposite to pure oak forest stands, may amplify Phytophthora inoculum in declining oak plots. By contrast, in non-declining oaks, all species of *Phytophthora* were more likely to be abundant in the rhizosphere of oaks colonised by *Bacillus* when stands were dominated by oaks. Furthermore, the colonisation of roots by the mold *Penicillium* and the Actinomycete *Acidothermus* were negatively correlated with *Phytophthora* occurrence. Such correlations may reflect shared environmental/habitat preferences or interactions between the microorganisms. Several studies on cultivated plants, infected or not by diverse *Phytophthora* species, have revealed that the root colonization by pathogens induce modifications of the root microbiota (Larousse and Galiana 2017, Vlassalaer et al 2024). Conversely, members of the root microbiota can reduce or stimulate the colonisation by *Phytophthora. Bacillus, Penicillium* and Actinomycetes regroup many strains well known for their ability to produce secondary metabolites including active metabolites against diverse species of *Phytophthora* and some species are used as biocontrol agents against root pathogens (Van Loocke etal. 2025). By contrast, *B. megaterium* Sb5 was found to stimulate the infection of soybean by several species of *Phytophthora* (Kong & Hong, 2016). More data are needed to complete and validate the predictive models obtained here and to determine how *Phytophthora* abundance in oak roots is influenced by the local microbiota and *vice versa*. However, our results suggest that monitoring fungal and bacterial microbiota, together with tree species diversity, may enable the prediction of susceptibility to *Phytophthora* host colonisation. The identification of microbial biomarkers of susceptibility could be implemented in plant production in nurseries and breeding programs for reforestation. Microbial biomarkers can potentially improve forest monitoring as early warning signals of forest decline, to help anticipate future crises.

### Combination of fungal and bacterial ASV best predicted oak health status

*Phytophthora* DNA occurrence in the bulk soil, rhizosphere or roots of oaks was not a good predictor of the health status of the trees. Instead, we obtained a model explaining well the measured variability by combining fungal and bacterial variables: the relative abundance of two BS bacterial ASVs, one R bacterial ASV and the proportion of high biomass ECM exploration type. Although this model requires validation using larger, independent datasets, to the best of our knowledge, this is the first study to use non-pathogenic microbiota to predict oak health status. These findings highlight the value of multi-kingdom approaches and holobiont-level studies for developing a more comprehensive understanding of the decline process. Using also a multi-kingdom approach, Koskella and colleagues showed that microbial communities of the bark of chestnut trees confer benefits to healthy trees and reduce the risk of diseases (Koskella et al. 2017, Feau & Hamelin 2017). Machine learning and statistical models are increasingly used to predict phenotypic or health outcomes based on microbiome data, mainly in the human health field (Marcos-Zambrano et al. 2021) but we can expect that such studies will be on the rise in the forest field.

### Oak decline locally affected microbial functional activities

The Chantilly forest spreads over a mosaic of soils ranging from acidic and nutrient-poor podzols to nutrient-rich and fertile luvisol (Saintenoise et al. 2024). As expected (Uroz et al. 2016, Baldrian 2023), this strongly drives the composition of soil and oak-associated fungal and bacterial communities. Despite this spatial heterogeneity of community composition, most enzymatic activities involved in C, N and P cycling measured in the rhizosphere of healthy oaks were similar across soil types suggesting functional redundancies among the main players of nutrient cycling in the soil, in accordance with previous reports (Colin et al. 2017, Zhang et al. 2016, Zeng et al. 2023, Auer et al. 2024). Two enzymatic activities involved in C cycling (chitinase and laccase) were consistently decreased in the rhizosphere of declining oaks but only on specific soil types. Chitinases participate in the degradation of chitin, the main component of cell walls of fungi and insects and thus may be linked to a reduction of fungal biomass (Baldrian et al. 2023). Conversely, laccases are produced by fungi to degrade lignin in search for N in recalcitrant plant organic matter. Surprisingly, while this activity strongly correlated with the ECM *Russula* in the rhizosphere of non-declining oak, such correlation was not observed in declining oak samples. This may suggest that *Russula*, despite still colonizing oak roots, would have a reduced laccase activity when oaks are under decline. However, the main actors of organic matter degradation in forest soils are the saprophytic fungi. Therefore, the reduction of laccase activity could also be due to a reduction of the activities of saprophytic fungi despite their abundance remaining unchanged. Further studies are needed to determine the mechanisms behind enzymatic reduction, but these results suggest that the functional activities of microbial communities can be modified in response to oak decline, even if their composition remains unchanged.

## CONCLUSION

Data acquired in the frame of this study suggest that oak-associated microorganisms are not responsible for oak decline in the Chantilly forest but that the alteration of the microbial communities may contribute to oak weakening through a combination of effects: a higher colonisation by pathogenic *Phytophthora*, the reduction of the association with high biomass ECM fungi with long distance exploration abilities, and the reduction of the ability to recycle organic matter in specific soil types. While oak decline seemed to impact the colonisation of roots by specific fungal and oomycetes taxa, bacterial response was more sensitive in the bulk soil, revealing potential indirect effects of decline which would deserve to be further explored. Furthermore, integrative multi-kingdom analyses indicate correlations between the abundance of *Phytophthora* species and certain bacterial and fungal taxa which may favour or reduce the colonisation by weakening pathogens. The extension of such a type of study at a larger scale could likely permit to predict microbial patterns, soil conditions and forest management practices which enhance the risk of oak decline.

## ACKNOWLEDGEMENTS

Acknowledgements are expressed M. Bednareck, T. Roussel, the association “Sauvons la forêt de Chantilly”, the staff of ONF and Institut de France for their help in the sample collection. We are grateful to N. Hirt, R. Charrier (BEF INRAE) and the members of volunteers of the collective “Sauvons la forêt de Chantilly” and the ONF team for the establishment of permanent survey plots and data acquisition, N. Leroy (SILVA INRAE) for her assistance in using QGIS, B. Bracchi (BIOGECO, INRAE) for providing information regarding tree genotypes, and the PGTB INRAE platform for providing help during library construction and sequencing processes. Lastly, we would like to thank the Institut de France and the Château de Chantilly for giving access to the forest site and actively supporting this research.

