## Supplemental Information for "Microbial communities do not drive chronic *Quercus* spp. decline but exacerbate tree weakening: a multi-kingdom soil–root continuum study in the Chantilly forest"

**Table of contents**

1. Chantilly forest site description
2. Sample site properties
3. Supplemental figures
4. References

#### 1. Chantilly forest site description

Chantilly forest is situated in the Oise department in France, 40 km North East of Paris. It spreads over 6 300 ha and it is mainly located on a Lutetian limestone plateau covered with Aeolian sands. The stands are mainly made up of an aged, open grove of pedunculate and sessile oaks on an aged coppice dominated by small-leaved lime. *Quercus* species make up 48% of the forest, with an average age of 150 years and a low density of 35 oaks/ha.

Oak dieback has been reported in this forest for around twenty years, and the phenomenon accelerated towards 2017. Given this situation, a joint participatory research project involving managers, the public and researchers aiming at monitoring and understanding process of die back has been set up in 2018. In this frame, two sets of plots have been defined : a set of 300 plots was first used to monitor dieback level per plot according DEPERIS protocol (https://agriculture.gouv.fr/la-methode-deperis-comment-quantifier-et-mesurer-letat-de-sante-dune-foret-et-son-evolution). A subset of 60 permanent plots (2 x 30 meters in diameter) were then established among the 300 plots for in-depth description and monitoring. The 22 plots used in this study are part of the 60 permanent plots further described. In 2021, pedologic pits from 110 to 230 cm were dug to determine qualitative (soil typology, humus) and quantitative variables (water pH, physicochemical measurements – total carbon and nitrogen stocks) which characterized the plot's soil. Circumference of each oak tree from every plot were measured and the proportion of fine roots, total biomass and the depth of roots was estimated for each plot. All metadata associated with the 60 permanent plots are available on the French dataverse (Saint-André et al. 2024[a,b] , Santenoise et al. 2024, Sénéchal-Chevalier et al. 2024). In 2021, a crown deterioration score (from A to F) was evaluated for each oak tree, based on the percentage of branch loss and branch mortality observed in the functional crown (part of the crown outside the tree's competition), according to the DEPERIS protocol (see table below).

**Table 1.** DEPERIS protocol for the estimation of functional crown of the tree.

| | | Branching loss | | | | | |
|---|---|---|---|---|---|---|---|
| | | 0 to 5% | 6 to 25% | 26 to 50% | 51 to 75% | 76 to 95% | 96 to 100% |
| **Branch mortality** | 0 to 5% | A | B | C | D | E | F |
| | 6 to 25% | B | B | C | D | E | F |
| | 26 to 50% | C | C | D | D | E | F |
| | 51 to 75% | D | D | D | E | F | F |
| | 76 to 95% | E | E | E | F | F | F |
| | 96 to 100% | F | F | F | F | F | F |

#### 2. Sample site properties

Physico-chemical properties and species composition of forest stands being known to influence the composition of forest microbial communities (Llado et al. 2018, Uroz et al, 2016), we tested which variables separated the 22 plots in which the 29 pairs were sampled (see **Figure 1** below). While plots were not clearly separated by soil types, as illustrated by the PCA plot, forest management and tree characteristics (proportion of *Q. robus* vs *petrea,* proportion *Quercus* vs other species in the stand, tree circumference) explained a third of the variance while a second third was explained by physico-chemical soil properties (mainly pH and nitrogen-related variables). However, strong local variations were observed (Saintenoise et al. 2024). No correlation between variables was found (Pearson correlation $p>0.01$), thus these different variables were further used to test their influences on microbial community composition and activities.

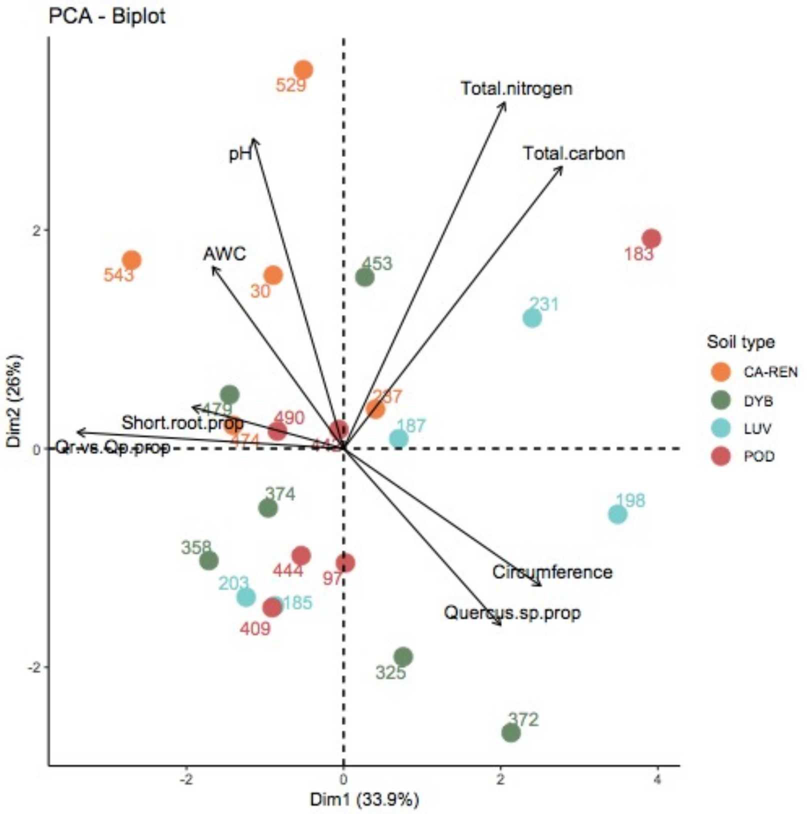


**Figure 1**. Principal Componant Analysis biplot showing distribution of the sampled plots depending on soil physico-chemical properties and tree species and tree properties: AWC – Available Water Capacity, Total nitrogen (%), Total Carbon (%), Circumference -diamater at breast hight, Quercus sp. prop – proportion of Quercus species in the plot, Qr vs Qp prop – proportion of *Q. robur* vs *Q. petrea* in the plot, short root prop – proportion fo short roots vs bigger roots in soil. Each plot is represented by a point associated with its plot number and color coded based on soil typology: orange CA-REN, green DYB, blue LUV, red POD.

#### 3. Supplemental figures

##### Supplemental Figure 1

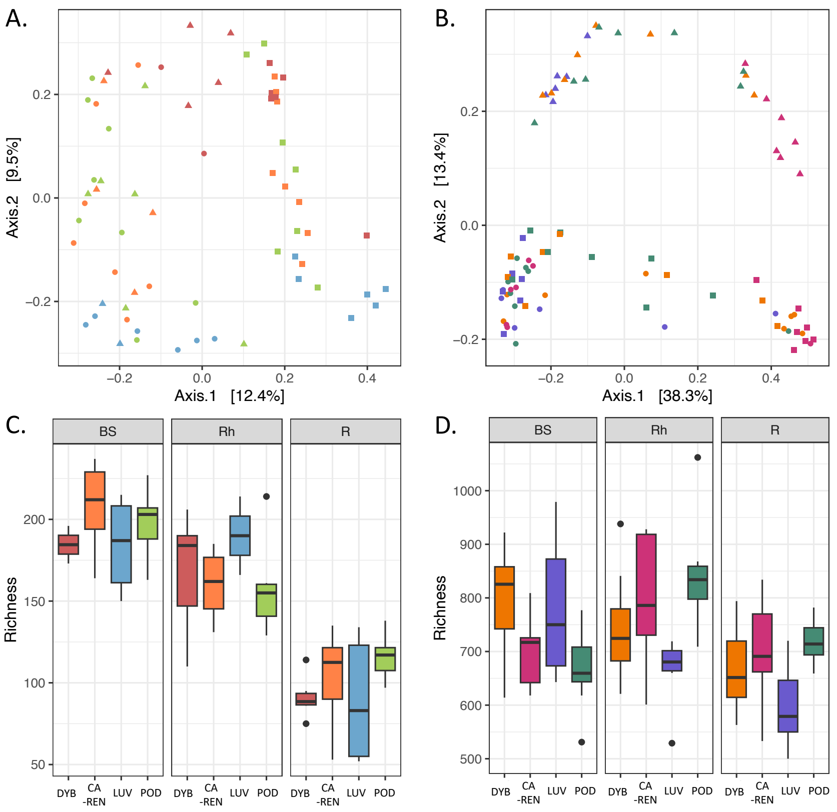


**Suppl. Figure 1. Microbial community structures and diversities of non-declining oaks as a function of compartment and plot soil type**. Principal Coordinate Analysis based on Bray-curtis dissimilarity matrices fungal (A) and bacterial (B) communities of non-declining. Each dot represents a sample, different compartment are represented by different shapes: dot – bulk soil, square – rhizosphere, triangle – root, different soil types are represented by different colors: orange- DYB, red – CA-REN , blue – LUV, green – POD. Richness as defined by total number of fungal (C) and bacterial (D) ASVs.

**Supplemental Figure 2**

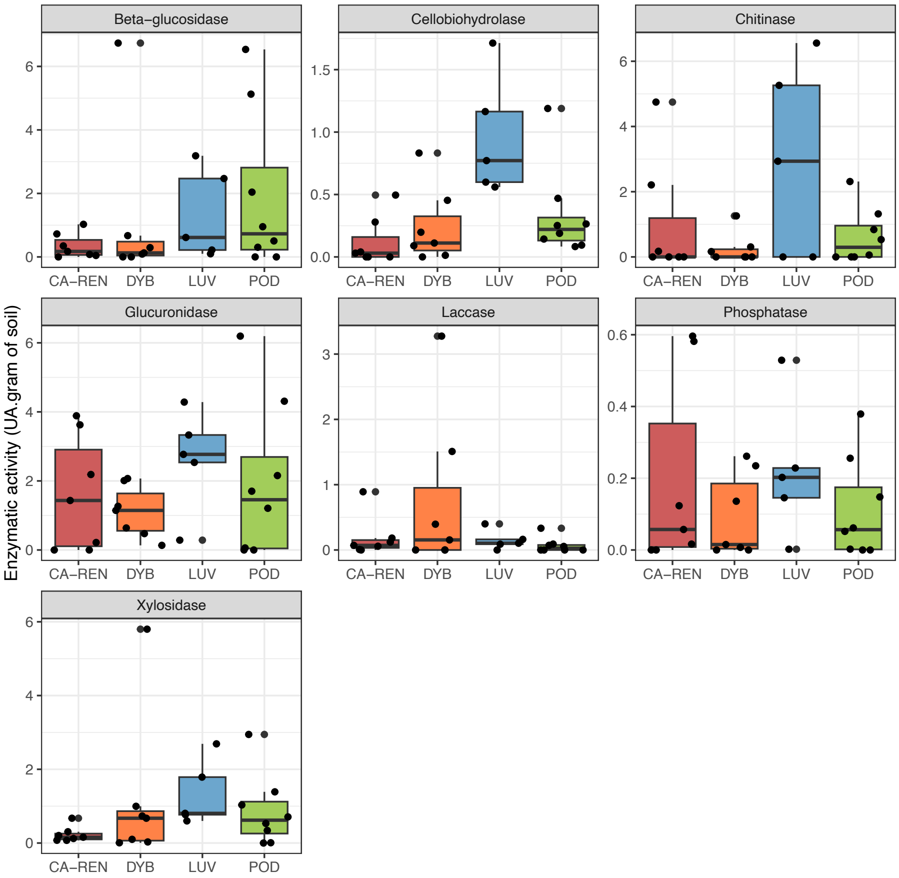


**Suppl Figure 2**. **Microbial enzymatic activities in rhizospheric soil of non-declining oaks as a function of soil types.** Generalized mixed model using sanitary status x soil type as fixed effects and pair as random effects with a Gaussian distribution was used to test the combined effects soil type on enzymatic activities. BH post-hoc test was used to correct for multiple testing. Corrected p-values ≤ 0.01 were considered significant

### Supplemental Figure 3

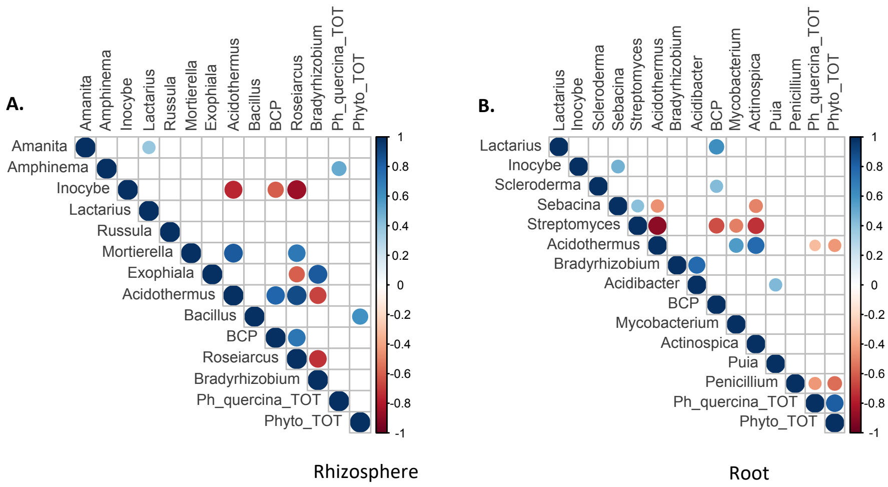


**Suppl. Figure 3**. Links between *Phytophthora* abundance and the relative abundance of other microbial taxa according to habitats. A. B. Spearmen correlation matrices between the relative abundance of the 12 most abundant fungal and bacterial taxa and the relative abundance of *Phytphthora quercina* and of all *Phyophthora* detected in the rhizosphere (BA) and the roots (B) of non declining oaks. Only statistically significant correlations are depicted (*p*. value <0.01). BCP = *Burkholderia-Caballeronia – Paraburkholderia*. Phyto_TOT = sum of all reads of all *Phytophthora* detected.

#### Supplemental Figure 4

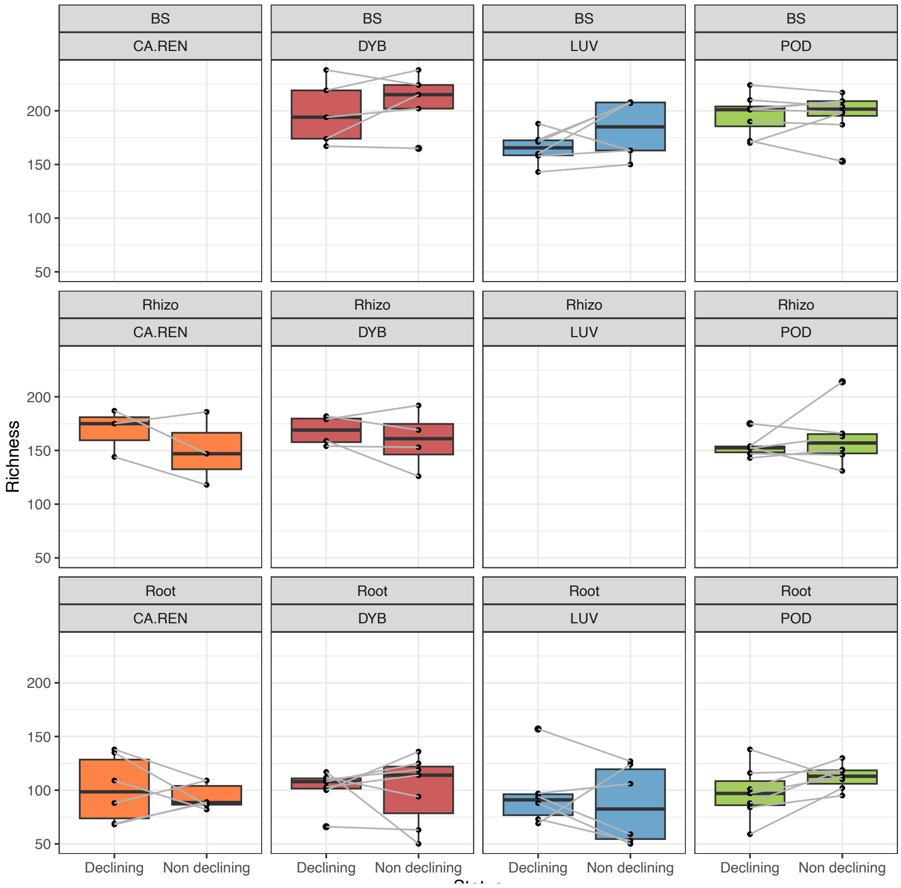


**Suppl. Figure 4**. Fungal richness of non-declining vs declining oaks as a function of habitat and plot soil type.
